# Whole-genome sequencing analysis of wild-caught house mice *Mus musculus* from Madagascar

**DOI:** 10.1101/2021.09.10.459745

**Authors:** Kazumichi Fujiwara, Marie C Ranorosoa, Satoshi D Ohdachi, Satoru Arai, Yuki Sakuma, Hitoshi Suzuki, Naoki Osada

## Abstract

In Madagascar, the house mouse (*Mus musculus*) is thought to have colonized with human activities and is now one of the most abundant rodents on the island. In this study, we determined the whole-genome sequences of five Madagascar house mice captured from the wild. We examined the evolutionary history of the population by analyzing the mitochondrial and autosomal genomes. We confirmed that the mitochondrial genome lineages of Madagascar house mice formed a monophyletic clade placed at one of the most basal positions in the species. An analysis of autosomal genomic sequences indicates that the Madagascar house mice are genetically members of *M. m. castaneus* (CAS), but also contain genetic elements of *M. m. domesticus* (DOM) resulting from hybridization between subspecies. The signature of a strong population bottleneck 1000–3000 years ago was observed in both mitochondrial and autosomal genomic data. All samples showed strong genetic affinity to many CAS samples across a wide range of Indian Ocean coastal and island regions, with divergence time estimated around 4000 years ago. These findings support that the Madagascar house mice started to colonize the island with human agricultural activity, and experienced complex history for the establishment.

## Introduction

The house mouse (*Mus musculus*) originated from the northern part of the Indian subcontinent (Boursot *et al*., 1996; Din *et al*., 1996) and is one of the rodents that has experienced explosive population size growth and successfully colonized a wide range of continents, large islands, and even small remote islands around the world. Three major subspecies of house mice are considered to be inhabited around the world (Didion and Villena, 2013); the South Asian subspecies (*M. m. castaneus*: CAS), the North Eurasian subspecies (*M. m. musculus*: MUS), and the West European subspecies (*M. m. domesticus*: DOM). The house mice are small commensal mammals that spread worldwide along with the development of human agriculture. Therefore, analyzing colonization patterns of house mice would provide insight into the history of human migration (Prager *et al*., 1998; Duplantier *et al*., 2002; Searle *et al*., 2009a; Suzuki *et al*., 2013, 2015; Jing *et al*., 2014; Phifer-Rixey *et al*., 2018; Li *et al*., 2021, Britton-Davidian et al. 2007).

Madagascar is the fourth largest island in the world. It separated from the African continent around 160 million years ago, and also separated from the Indian subcontinent around 80 million years ago (Torsvik *et al*., 2000). The island is now about 300 km from the coast of the African continent at its shortest distance. Inferences from linguistic (Dahl, 1951, 1988; Adelaar, 1995), archaeological (Dewar and Wright, 1993; Burney *et al*., 2004), and genetic (Hurles *et al*., 2005) evidence suggest that humans in Madagascar are originated from both East Africa and Southeast Asia (in particular, Borneo Island). The question of when humans first arrived on the island of Madagascar is still an open and controversial issue. Although previous archaeological evidence has suggested that humans first settled on the island 1,500–2,000 years ago (MacPhee and Burney, 1991; Burney *et al*., 2004, Gommery et al. 2011), recent excavations have provided evidence that humans have been active on the island since even earlier than previously thought (Dewar *et al*., 2013; Hansford *et al*., 2018; Douglass *et al*., 2019). More recently, studies using human genome data showed that they migrated from Borneo approximately 1000–3000 years ago and from the east coast of Africa around 700–1500 years ago (Brucato *et al*., 2016 and Pierron *et al*., 2017). Subsequent intermittent human settlement on the island of Madagascar led to the formation of a large human population on the entire island approximately 1000 years ago (Battistini and Verin, 1972; Burney *et al*., 2004). At that time, a large commercial network connecting Asia and the Mediterranean Sea had been established along the coastal regions of the Indian Ocean (Verin and Wright, 1999), and traders from the Arabian Peninsula also sailed to the coast of Africa, including Madagascar (Allibert, 1988; Liszkowski, 2000).

Rodents have successfully colonized Madagascar island, and there are at least 23 species of rodents currently inhabiting it today (Rakotondravony and Randrianjafy, 1998). House mice are found in human settlements all over the island, though it is less abundant than the black rats (*Rattus rattus*), which are the most abundant rodent on the island (Goodman, 1995). In previous studies analyzing partial regions of mitochondrial genomes (control regions, the cytochrome *b* gene, and tRNAs), Madagascar house mice were genetically similar to samples from Yemen, in the southern part of the Arabian Peninsula (Duplantier *et al*., 2002; Sakuma *et al*., 2016). The mtDNA analysis revealed that the Yemeni house mouse mtDNA lineage forms a cluster of another potential subspecies, *M. m. gentilulus* (GEN), which is distinct from the three major subspecies (Prager *et al*., 1998; Suzuki *et al*., 2013). The mtDNA lineage in Madagascar constitutes a “narrow” monophyletic group, suggesting a recent and probably single origin (Duplantier *et al*., 2002; Sakuma *et al*., 2016). Suzuki et al. (2013) further compared the mitochondrial sequence of Yemeni GEN (from Prager et al. 1998) with the wild house mouse that Suzuki et al. had collected over a wide area of Eurasia, and the results showed that the mitochondrial lineage of the Yemeni GEN (which is again similar to the mitochondrial genome as Madagascar house mice) was in the most basal position to *M. musculus*. Note that it is unclear to which subspecies this Yemeni GEN belongs in terms of nuclear genes. In contrast, a microsatellite study suggested that Madagascar mice are genetically CAS (Hardouin *et al*., 2015). Furthermore, the phenotype of tail length is a typical character of DOM (Rakotondravony and Randrianjafy, 1998; Orsini *et al*., 1983; Marshall, 1998). These incongruences make the origin of Madagascar house mice highly controversial.

Previous studies on the genetic background of the house mouse in Madagascar are primarily based on mtDNA with limited nuclear genome analysis. In order to increase the resolution of genetic analyses, we performed high-quality whole-genome sequencing of five wild-caught *M. musculus* samples from the specific point from Madagascar and combined the data with previously published worldwide house mouse genomic dataset (Harr *et al*., 2016; Fujiwara *et al*., 2021). We determined the genetic position of these Madagascar wild house mice within the global population of *M. musculus* and estimated the divergence time to determine their genetic backgrounds and population dynamics. We consider that our study has provided new insights into the prehistoric migration history of house mice and humans in Madagascar.

## Materials and Methods

### Materials

Five specimens of the Madagascar wild house mouse were collected from the Parc Botanique et Zoologique de Tsimbazaza, Madagascar, and nearby areas which were also analyzed partial mitochondrial sequence and coat-color-controlling genes in the study by Sakuma et al. 2016 (Sakuma *et al*., 2016). The other genomic sequence data of *M. musculus* were obtained from previous studies (Harr *et al*., 2016; Fujiwara *et al*., 2021; Li *et al*., 2021). We used seven samples of the western Mediterranean mouse (*Mus spretus*) obtained from publicly available data (Harr *et al*., 2016). These data were downloaded from the DDBJ (PRJDB11027) and the European Nucleotide Archive (PRJEB9450, PRJEB11742, PRJEB14167, PRJEB2176, and PRJEB11897), representing the data of Fujiwara et al. (2021) and Harr et al. (2016), respectively. The *M. spretus* was used as an outgroup because it is a close relative to *M. musculus* and high-quality sequence data were publicly available for both the nuclear and mitochondrial genomes. The GRCm38 (mm10) was used as the reference genome sequence for *M. musculus*. The reference complete mitochondrial sequence of *M. spretus* (NC_025952) was also used for the mitochondrial lineage analysis. See Supplementary Table S1 for detailed information on the samples used in this study.

### Mapping genomic reads and variant calling

For the five Madagascar wild house mouse samples, 150-bp-length paired-end reads were generated using the DNBSEQ platform. The cleaned reads were quality checked using the FastQC v.0.11.9 tool (Andrews, 2010). The all cleaned reads were mapped to the house mouse reference sequence GRCm38 (mm10) using bwa-mem v.0.7.17-r1188 (Li and Durbin, 2009) algorithm with “-M” option and piped to Samblaster v.0.1.26 (Faust and Hall, 2014) program with “-M” option to mark PCR duplicated reads for downstream analysis. On the downloaded data from Harr et al. (2016), we used the samples that had greater than 20-fold median coverage. We used the GATK4 HaplotypeCaller (McKenna *et al*., 2010) function with “-ERC GVCF” to perform variant calls for single nucleotide variants (SNVs) in our dataset. The genomic variant call format (.gvcf) files of each sample were then merged to one jointly called file using GATK4 GenomicDBImport and GenotypeGVCFs functions. We then conducted the GATK4 Variant Quality Score Recalibration (VQSR) process to determine whether our newly sequenced raw variants are true positives or false positives, according to the machine learning method. For VQSR, we downloaded the known SNV dataset of “mgp.v3.snps.rsIDdbSNPv137.vcf.gz” from the Sanger Institute web server (ftp://ftp-mouse.sanger.ac.uk/REL-1303-SNPs_Indels-GRCm38/) as a training dataset. The HARD filtered SNVs with parameters “QD < 2.0, FS > 60.0, MQ < 40.0, MQRankSum < −12.5, and ReadPosRankSum < −8.0” was also used for training. After VQSR, we determined to use variants that passed within the 90% tranche (90% acceptance in all reliable training SNVs datasets) for downstream analyses. The autosomal SNVs were then filtered using genomic mappability scores to restrict our analyses to non-repetitive regions. To calculate mappability scores of the reference house mouse sequence, we run GenMap v.1.3.0 software with options “-K 30 and -E 2” and used the positions with the score of 1 (the 30-mer starting from the position is unique in the genome even allowing 2 mismatches) in downstream analyses. All samples were examined for kinship using the KING v.2.2.6 software (Manichaikul *et al*., 2010) with the option “--kinship.” None of the samples from Madagascar showed a kinship relationship. Therefore, a total of 133 samples of mice, five novel samples from Madagascar, and 128 publicly available samples were used in the following analyses.

Because the phenotypic records for sexes of Madagascar samples were not complete and may not be accurate, we determined the sexes of samples using the genomic sequence data. We calculated the read coverages on the sex chromosomes. We used the “depth” command of the samtools program to count the coverage of each sample in non-pseudoautosomal sites of the X and Y chromosomes that passed the mappability filter. The ratio of X to Y chromosome coverages exhibited a clear bimodal distribution in which the modes were 1.03–1.09 and 148.24–268.16. Three out of five samples with the higher ratios were judged as male samples (Supplementary Table S2). The synonymous and nonsynonymous SNVs were annotated with the house mouse gene annotation data version GRCm38.101 (ftp://ftp.ensembl.org/pub/release-101/gtf/mus_musculus/) using the SnpEff and SnpSift programs (Cingolani, Patel, *et al*., 2012; Cingolani, Platts, *et al*., 2012).

### Mitochondrial lineage analysis

The mitochondrial genome sequences of Madagascar mouse samples were reconstructed by a *de-novo* assembly software, Novoplasty v.4.2.1 (Dierckxsens *et al*., 2017), using the house mouse mitochondrial genome NC_005089 as a seed sequence. For mitochondrial genome analysis, *M. spretus* was used for the outgroup. The mitochondrial sequences of all samples were aligned using MUSCLE (Edgar, 2004a, 2004b) implemented in MEGA7 (Kumar *et al*., 2016), and all D-Loop regions and gapped sites were removed, resulting in 16,038 bp length. To construct the maximum likelihood (ML) tree of the mitochondrial genome, we used IQ-TREE v.1.6.12 (Nguyen *et al*., 2015). We also estimated the best substitution model using the ModelFinder (Kalyaanamoorthy *et al*., 2017) function implemented in IQ-TREE and used the “TIM2+F+R3” model for our calculations. The bootstrapping values were computed with 1000 replications. We used the program, BEAST v1.8.4 (Drummond and Rambaut, 2007) to estimate the time to most recent common ancestors (tMRCA) using mitogenome sequences (15,181 bp length excluding control region), the HKY+G substitution model, and the strict clock model as reported previously (Li *et al*., 2021). The Markov chain Monte Carlo simulation was run for 10 million steps (burn-in 10%) and sampled every 10,000 steps. Tracer v1.6 software (Rambaut *et al*., 2018) was used to assess the convergence of the chains and ensured effective sample size (ESS) values above 200 for most parameters. The trees were summarized using TreeAnnotator v1.8.4 software (http://beast.community/treeannotator) with the settings “Maximum clade credibility tree” and “Mean heights” and were displayed using FigTree v1.4.3 software (http://tree.bio.ed.ac.uk/software/figtree/). The time-dependent evolutionary rates of mtDNA (Ho *et al*., 2011) were considered as previously described (Li *et al*., 2021). The evolutionary rates of 2.4 × 10^−8^ substitutions/site/year and 1.1 × 10^−7^ substitutions/site/year were used for the older (> 100,000 years) and younger divergences (< 20,000 years ago), respectively. For the results output from the BEAST analysis, fixing the divergence time scale was performed by using the time-dependent evolutionary rate described above.

### Population structure analysis of the Madagascar house mouse

We performed SNV filtering by using vcftools omitting indels and multiallelic SNVs for downstream analysis. The filtered autosomal SNVs were then converted to a PLINK v.1.9 (Purcell *et al*., 2007) format. In general practice, the SNVs in linkage disequilibrium (LD) are excluded from the population structure analysis. The house mouse samples were genetically highly structured at the subspecies level and excluding these SNVs would result in a very small number of SNVs left for analysis. We performed principal component analysis (PCA) and Admixture analysis both with and without LD pruning.

PCA was performed using the smartpca program implemented in Eigensoft v.7.2.1 (Patterson *et al*., 2006). In order to calculate eigenvalues, the default parameter settings for smartpca were used, except that we did not remove outlier samples.

We computed *f*_3_-statistics and *f*_4_-statistic because both statistical methods are very useful statistics to infer the relationship among populations. Both methods were introduced by Patterson et al. 2012, measuring allele frequency correlations among selected populations (Patterson *et al*., 2012). The *f*_4_-statistics were computed using the qpDstat program with “f4 mode” and “printsd” options implemented in AdmixTools v.7.0 (Patterson *et al*., 2012). The Indian CAS and German DOM samples were used as reference populations as CAS and DOM. respectively. These samples showed the least amount of admixture with other subspecies based on the *f*_4_ statistics in the previous study (Fujiwara *et al*., 2021). The configuration of the *f*_4_-statistics is represented as *f*_4_(A, B; C, D), where A to D represents each population. If the value of *f*_4_(A, B; C, D) is not statistically significantly different from 0, the allele frequency difference between A and B would be independent on the allele frequency difference between C and D. In contrast, if the value of *f*_4_(A, B; C, D) is statistically significantly positive, the gene flow may exist between A and C or B and D. Furthermore, if the value of *f*_4_(A, B; C, D) is statistically significantly negative, the gene flow may exist between A and D or B and C. The outgroup *f*_3_ statistics were computed by the qp3pop program of AdmixTools using the “outgroupmode” option. In the outgroup *f*_3_-mode, the statistics of *f*_3_(A, B; C), where C represents an outgroup population, show the genetic drift from C to the common ancestor of A and B and indicate the genetic relatedness between the population A and B.

A neighbor-joining tree was reconstructed using a pairwise distance matrix generated from autosomal data (Saitou and Nei, 1987). Our previous study (Fujiwara et al., 2021) showed that samples with a high proportion of admixture may considerably distort the pattern of divergence among subspecies. For example, the inclusion of RUS01 from Moscow, which showed 56% DOM ancestry and 43% MUS ancestry in the results of ADMIXTURE v.1.3 (Alexander et al., 2009) analysis (K = 3), largely changed the blanching pattern of subspecies. Therefore, in this study, we performed ADMIXTURE analysis with K = 3 and constructed phylogenetic trees using only those samples in which ancestry to a particular subspecies accounted for more than 80%. We computed all pairwise identity-by-state (IBS) distances using the PRINK software with the “--distance square 1-ibs” option (Purcell *et al*., 2007). The tree was visualized using the Ape package in R (Paradis *et al*., 2004). The Neighbor-Net tree (Bryant and Moulton, 2004) was computed using the phangorn package (Schliep, 2011) with gdsfmt, SeqArray, SNPRelate packages (Zheng *et al*., 2012, 2017) in R.

### Analysis of genomic local ancestry

To simulate the local ancestry inference of our Madagascar samples, we applied the Loter software (Dias-Alves, et al. 2018) to five phased samples from Madagascar and four phased samples from Indonesia. For reference populations, we selected CAS from India and DOM from Germany as possible source populations, assuming that haplotypes of Madagascar and Indonesia samples consist of mosaic structures of these DOM and CAS samples.

### Demographic inference

Pairwise sequentially Markovian coalescent (PSMC) (Li and Durbin, 2011) analysis was performed for all five samples from Madagascar. To create the input file for PSMC, we used the “mpileup” command of samtools with the “-C 50, -O, -D 60, -d 10” option to obtain the consensus autosomal genome sequence with GRCm38 (mm10) as the reference genome. The PSMC analysis was conducted under the option with “-N25 -t15 -r5 -p ‘1*4+25*2+1*4+1*6’”. We performed 100 bootstrap replicates for each representative subspecies sample to test the variance of the estimated effective population size (*N*_e_). Generation time was assumed to be 1 year (Suzuki et al. 2004, Geraldes et al. 2008, and Bronson 1979), considering the mating in the wild environment, and a mutation rate of 0.57 × 10^−8^ per site per generation (Milholland *et al*., 2017).

The multiple sequentially Markovian coalescent (MSMC) (Schiffels and Durbin, 2014) and its second version (MSMC2: https://github.com/stschiff/msmc2) (Malaspinas *et al*., 2016; Schiffels and Wang, 2020) were used to estimate *N*_e_ changes and population separation history. In our MSMC/MSMC2 analysis, we performed estimations using a phased haplotype sequence as input data. For phasing, ShapeIt4 software (Delaneau *et al*., 2019) was used to generate phased haplotype data. We performed the phasing procedure to 126 house mouse samples used in this study; we did not include seven *M. spretus* samples for the phasing procedure since we do not have the recombination rate for them. The “Mapping Data for G2F1 Based Coordinates” (Liu *et al*., 2014) from “Mouse Map Converter (http://cgd.jax.org/mousemapconverter/)” was downloaded to provide the recombination rate input file for ShapeIt4 and also MSMC/MSMC2. On MSMC/MSMC2 analysis, the mappability score of the reference mouse genome is considered, and only unique sequence positions were used in the calculations. To estimate the population separation history for CAS and Madagascar, or DOM and Madagascar, we used two haplotypes from each population (four haplotypes total) to calculate the relative cross coalescent rate (rCCR) for given population pairs. According to Shiffels et al. (2014) (Schiffels and Durbin, 2014), the rCCR variable ranges between 0 and 1 (in some cases it is unavoidable to calculate greater than 1), and the value close to 1 indicates that two populations were not differentiated. Heuristically, the half value of rCCR (i.e., rCCR = 0.5) is assumed as the time when the two populations separated. In the bootstrapping method of MSMC2, the original input data was cut into blocks of 5Mbp each, which were then randomly sampled to create a 3Gbp-length pseudo genome. Calculations were performed on a total of 20 of these pseudo genomes. To visualize the MSMC/MSMC2 data, we assumed a generation time of 1 year (Suzuki et al. 2004, Geraldes et al. 2008, and Bronson 1979) and a mutation rate of 0.57 × 10^−8^ per site per generation (Milholland *et al*., 2017).

## Results

### *Genetic diversity of Madagascar* house mice

We analyzed five whole-genome sequences of *M. musculus* from Madagascar, partly analyzed in the previous study (Sakuma *et al*., 2016). These Madagascar samples were jointly genotyped with 128 samples, including seven *M. spretus* samples, which were obtained from the previous study (Fujiwara *et al*., 2021). Supplementary Table S1 contains a list of Madagascar samples with sample ID and approximate captured locations. The number of filtered SNVs in five Madagascar samples was 24,883,782 in total with an overall transition/transversion ratio of 2.10. The average per-sample nucleotide diversity (heterozygosity) of the Madagascar sample was 0.0031. Table 1 presents the detailed metrics for each Madagascar wild house mouse sample.

**Table 1.**
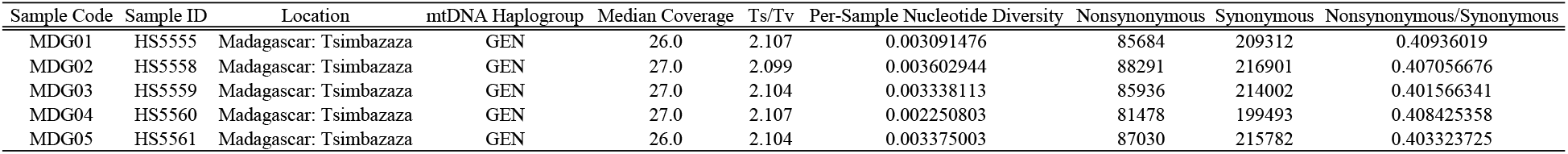
Basic Statistics of Malagasy House Mouse Samples.

### Population structure

To define the genetic features of the Madagascar house mice, we first performed a PCA using autosomal 86,288,314 SNVs from 126 *M. musculus* samples (Fig. 1). In the PCA plot, three clusters on the vertices of a triangle correspond to MUS, CAS, and DOM genetic components, and individuals lie on the intermediate shows inter-subspecies hybrid samples. The PC1 (eigenvector 1) indicates the genetic difference between MUS and CAS, and the PC2 (eigenvector 2) indicates the genetic difference between the DOM and CAS. The five Madagascar samples were positioned near the CAS cluster with a slightly admixed genetic component of the DOM. There was little variation in eigenvectors among the Madagascar samples. The LD pruned version of PCA was also analyzed, but the results were not significantly different from Fig. 1 (Supplementary Fig. S1). We also performed ADMIXTURE analysis to estimate ancestral genetic components in the Madagascar samples (Supplementary Fig. S2), assuming three (K = 3) and four (K = 4) ancestral populations. In both results, the Madagascar samples were found to have a large CAS ancestry and a small DOM ancestry (approximately 87% CAS ancestry).

**Fig. 1.**
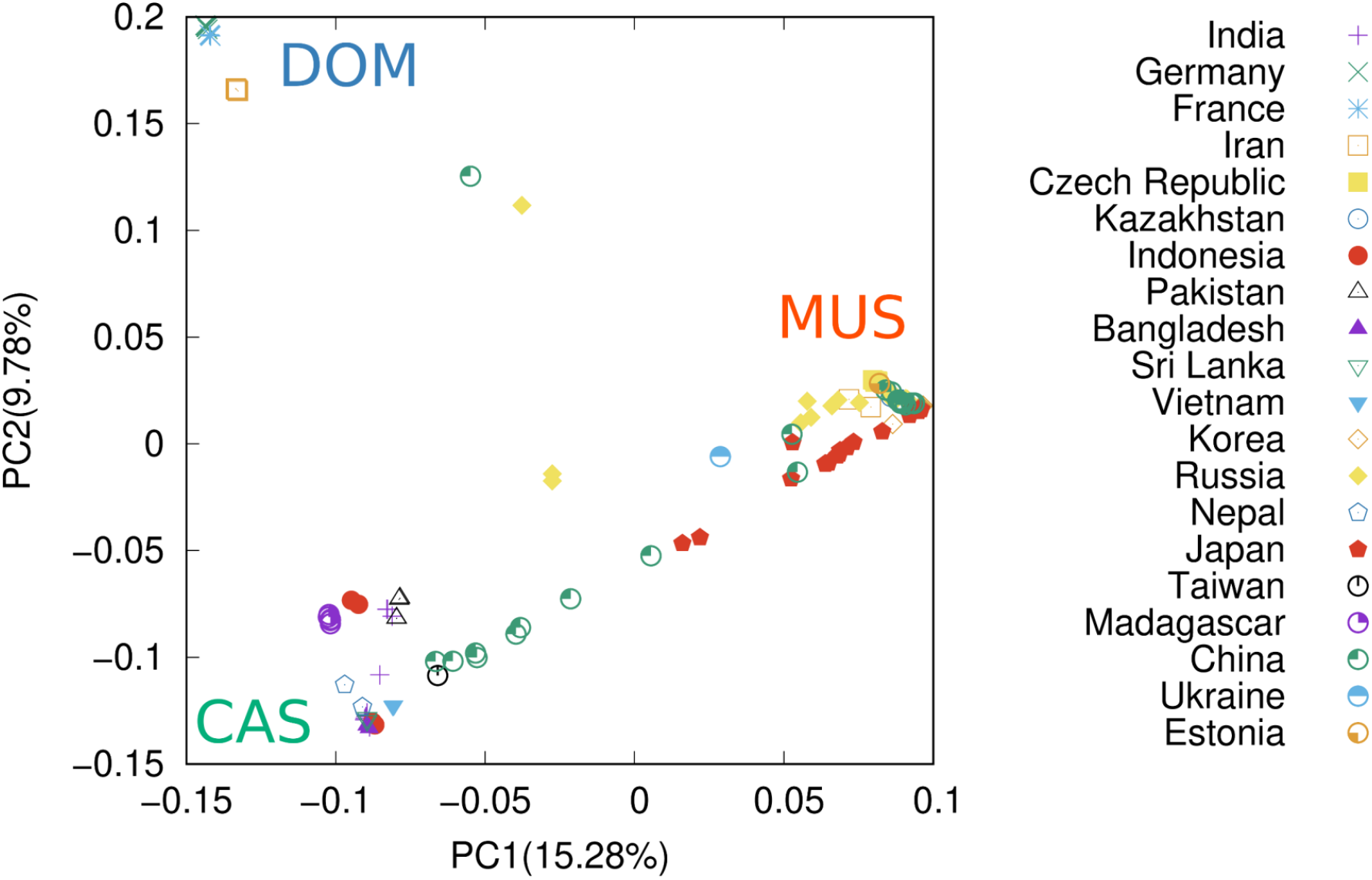
Principal Component Analysis plot of *M. musculus* using autosomal single nucleotide variants (SNVs). The points in the figure represent the eigenvalues of individual samples collected from the countries shown on the right. The upper left, lower left, and right vertices of the triangle represent *Mus musculus domesticus* (DOM), *M. m. castaneus* (CAS), and *M. m. musculus* (MUS) genetic components, respectively. The proportion of variance for each eigenvalue is shown in parentheses on the labels of the *x*-axis and *y*-axis. The arrow shows the position of Madagascar samples.

Next, we constructed an autosomal genetic tree using the neighbor-joining method with the *M. spretus* population set as the outgroup (Fig. 2). The samples with less than 80% of specific ancestral components in the ADMIXTURE analysis were not used in the phylogenetic analysis (see Methods for detail). The tree showed that the Madagascar samples formed a single cluster within the CAS cluster. Neighboring samples of the Madagascar samples were the Nepalese samples and the Indian samples except for the mountainous areas. Furthermore, we constructed a neighbor-net network (Supplementary Fig. S3). The neighbor-net network showed a similar pattern to the neighbor-joining tree, showing that the Madagascar wild house mouse samples are within the CAS samples clusters. To get a clearer picture, we used the outgroup *f*_3_ statistics, *f*_3_(MDG, X; SPR), where MDG, X, and SPR represent Madagascar, all non-Madagascar *M. musculus*, and *M. spretus* population, respectively. The results showed that individuals genetically close to the Madagascar samples (high *f*_3_ values) were located around the coastal side of the South Asian regions and Indonesian islands (Fig. 3).

**Fig. 2.**
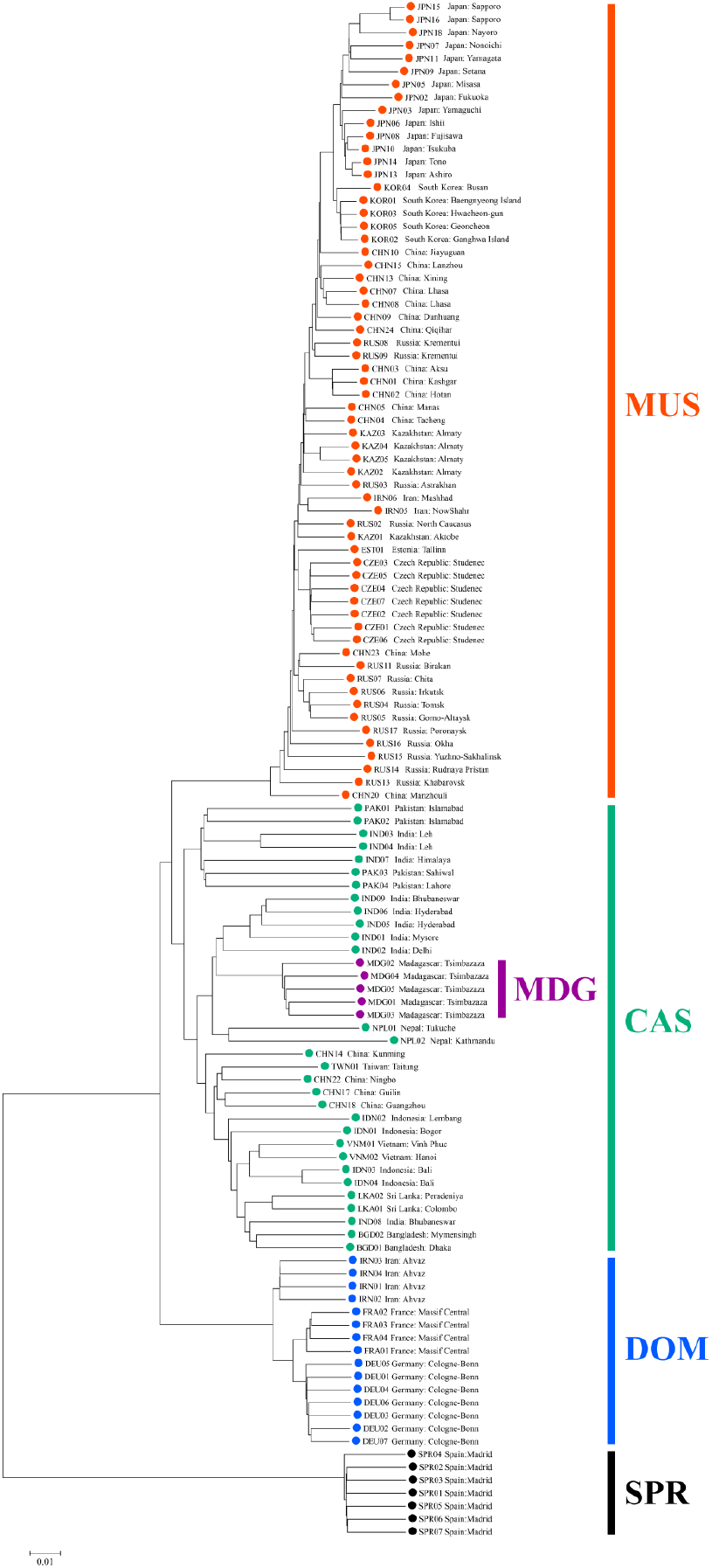
The neighbor-joining tree was inferred using pairwise genetic distances of *M. musculus*. The seven samples of *M. spretus* (SPR) were used as the outgroups. The color shows the subspecies: red (*M. m. musculus*: MUS), green (*M. m. castaneus*: CAS), blue (*M. m. domesticus*: DOM), and cyan (*M. musculus* - Madagascar). The hybrid samples were excluded from constructing the distance tree.

**Fig. 3.**
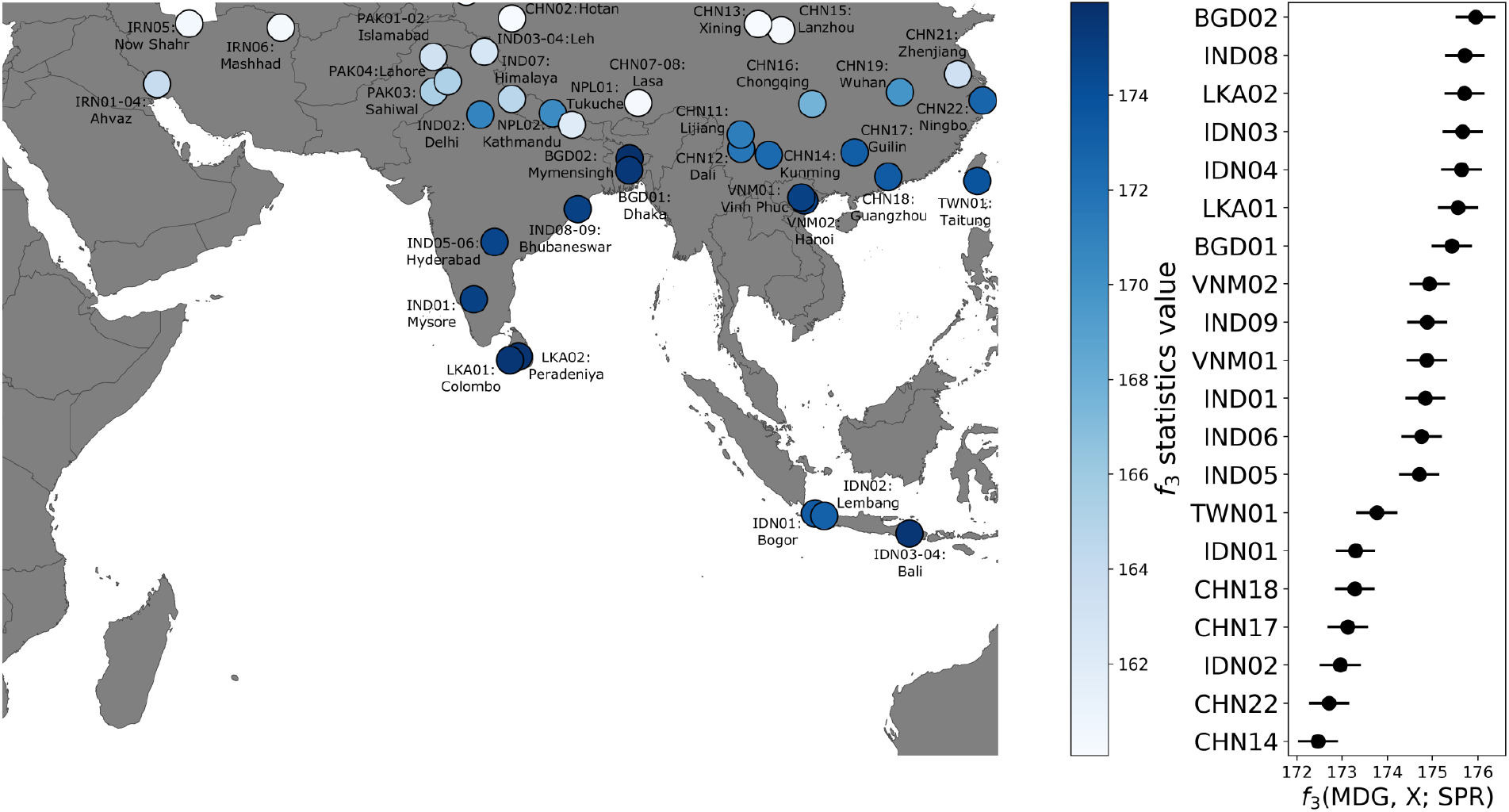
Outgroup *f*_3_ (MDG, X; SPR) statistics. The right plot shows the top 20 of the outgroup *f*_3_ statistics values in which individuals of regional origin are genetically close to the Madagascar wild house mice. The horizontal line associated with each point indicates the standard error. The higher values indicate genetic closeness to the Madagascar house mice. The left world map shows the *f*_3_ statistics value heat map focused on the South Asian region.

We investigated the admixture pattern of Madagascar samples using *f*_4_ statistics. Fujiwara et al. (2021) reported that Indian and German samples experienced the smallest amount of gene flow among subspecies and would be the reference populations for CAS and DOM, respectively. We used 102,858,288 SNVs to compute *f*_4_(SPR, CAS; DOM, MDG) and *f*_4_(SPR, DOM; CAS, MDG). In both cases, the *f*_4_-statistics scores indicate significantly positive values. The Z-score was 52.43 for *f*_4_(SPR, CAS; DOM, MDG) and 26.25 for *f*_4_(SPR, DOM; CAS, MDG), suggesting that the Madagascar samples were genetically closer to CAS than to DOM, with a non-negligible amount of gene flow with DOM. Supplementary Table S3 presents detailed results.

### Mitochondrial lineage analysis

We further clarified the genetic background of Madagascar samples using complete mitochondrial genomes. Phylogenetic trees based on the maximum likelihood of inference using the complete mitochondrial genome revealed that the five individuals from Madagascar formed a distinct cluster (Fig. 4a, Supplementary Fig. S4). The Madagascar samples were found in the most basal lineage of all *M. musculus* mitochondrial haplotypes, but bootstrap support was not sufficiently high (0.64, Supplementary Fig. S4). The tMRCA of Madagascar mitochondrial lineages was estimated using the BEAST software and the analysis showed that the expansion of Madagascar samples occurred about 3,000 years ago (Fig. 4b).

**Fig. 4.**
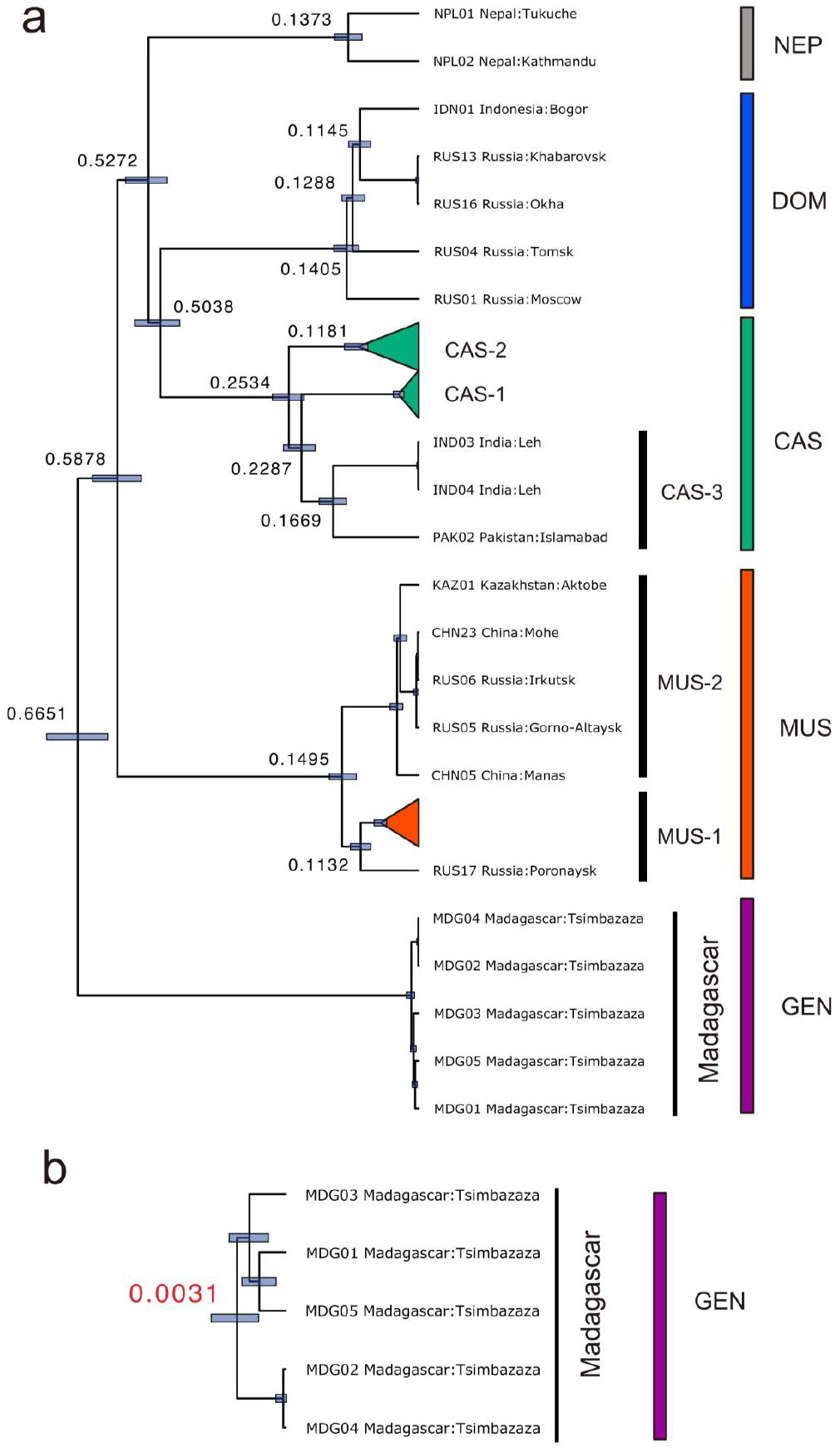
Divergence time estimates (million years ago) of the five haplogroups of the *M. musculus* subspecies based on entire mitochondrial genome sequences (15,000 bp) and a Bayesian-relaxed molecular clock of 2.4 × 10^−8^ substitutions/site/year (**a**). Blue bars represent the 95% highest posterior density interval. See Li et al. (2021) for the details of the subclades. Given the time-dependency of the mtDNA evolutionary rate, the meantime of the most recent common ancestor of the five haplotypes from Madagascar was estimated with a molecular clock of 1.1 × 10^−7^ substitutions/site/year (**b**).

### Inference of past demographic history

PSMC analysis was performed to estimate the demographic history of the Madagascar house mouse samples. The PSMC plot showed that all Madagascar samples experienced similar effective population size changes over the past 1,000,000 years (Fig. 5). The ancestral population experienced a rapid increase in effective population size from 200,000 to 300,000 years ago, peaked around 100,000 years ago, and has been slowly declining since then over the Last Glacial Period. The PSMC plots for the representative individuals of CAS and DOM are also shown in Fig. 5. for comparison. According to the PSMC plot, the DOM, CAS, and Madagascar populations experienced similar population histories until about 400,000–500,000 years ago, and the DOM populations experienced different population size trajectories from the other populations afterward. CAS and Madagascar populations experienced similar population histories until about 100,000 years ago. The CAS population has experienced a rapid decrease in effective population size after 100,000 years ago. The Madagascar samples also experienced a similar decline, but with a slower rate than the CAS population. We also performed MSMC analysis using 10 haplotypes from the Madagascar samples to analyze the relatively recent effective population size (Fig. 6). The plot exhibits a historical bottleneck event of the Madagascar samples around 1000–3000 years ago.

**Fig. 5.**
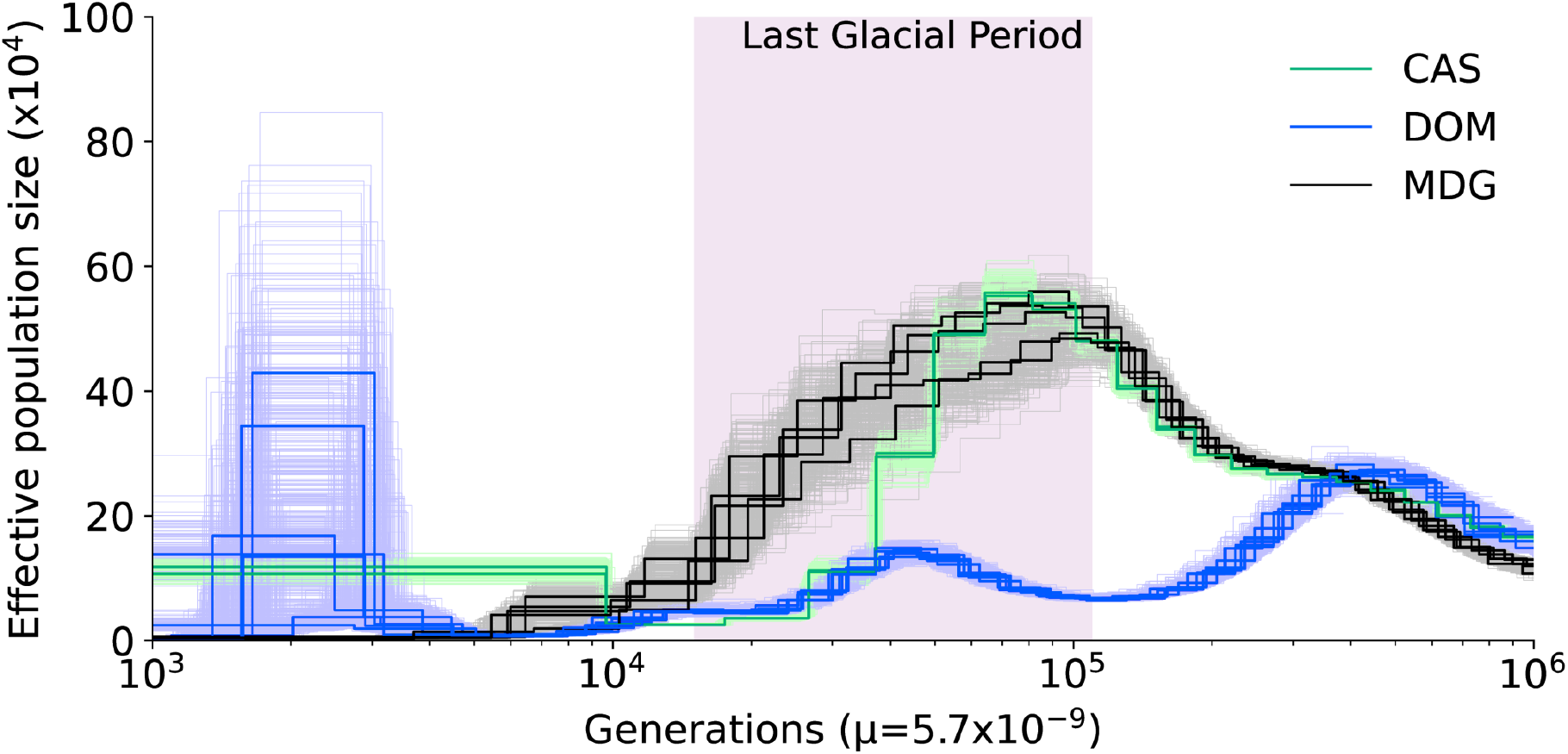
The Pairwise Sequentially Markovian Coalescent (PSMC) plot of *M. musculus*. The green, blue, and black lines represent the inferred effective population size transition of CAS, DOM, and Madagascar (MDG) mouse populations, respectively. The *x*-axis represents generations before the present scaled by the mutation rate 0.57 × 10^−8^ per site per generation and the y-axis represents the number of the inferred effective population size. The lines with lighter colors represent 100 replications of the bootstrapping results.

**Fig. 6.**
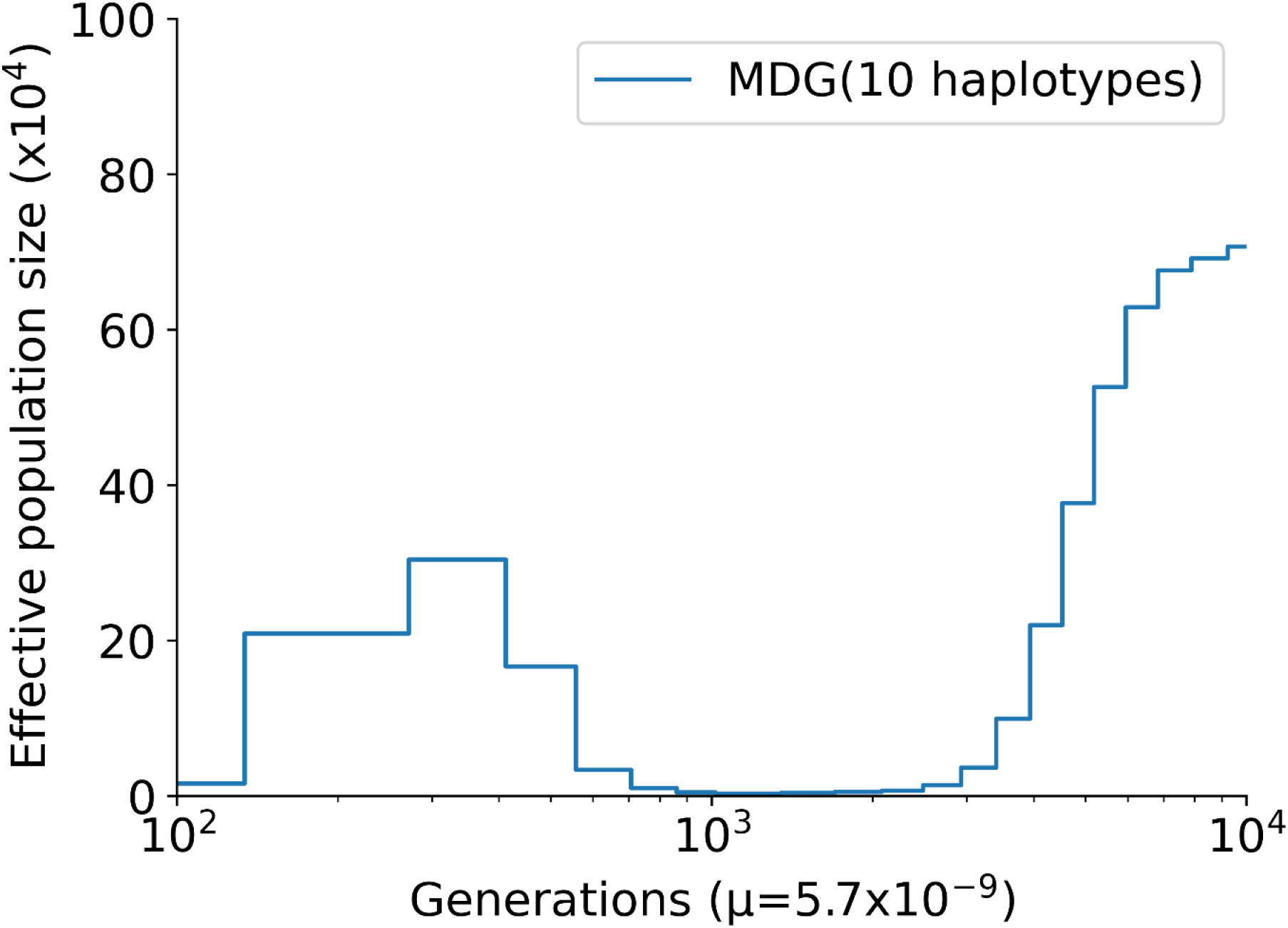
The Multiple Sequentially Markovian Coalescent plot of five Madagascar samples using 10 haplotypes. The *x*-axis represents generations before the present scaled by a mutation rate of 0.57 × 10^−8^ per site per generation. The *y*-axis represents the number of the inferred effective population size of Madagascar house mice.

### Estimation of divergence time between populations

In order to estimate when the Madagascar samples diverged from the other populations, we additionally performed the cross-population MSMC analysis. MSMC analysis was performed using two haplotypes for each sample. We evaluated cross coalescent rates between Madagascar samples and 13 different samples, including German DOM, Indian CAS, and 11 CAS individuals from the coastal and island regions of the Indian Ocean such as from India, Bangladesh, Sri Lanka, and Indonesia. The divergence time between German DOM and Madagascar, which was approximated with the time of rCCR = 0.5 (see Methods), was approximately 225,000 years ago with 95% Confidence Interval (CI) of 224,641–226,546. While an Indian CAS, which was sampled at the inland mountainous area, showed the divergence time of approximately 50,000 years ago (95% CI: 48,907– 50,011) from Madagascar, the other CAS samples from the coastal regions of the Indian Ocean showed the much more recent (∼4000 years ago) divergence time from Madagascar (Fig. 7). As a control, we calculated rCCR on two samples within Bali and found that the two samples did not diverge (Supplementary Fig. S5).

**Fig. 7.**
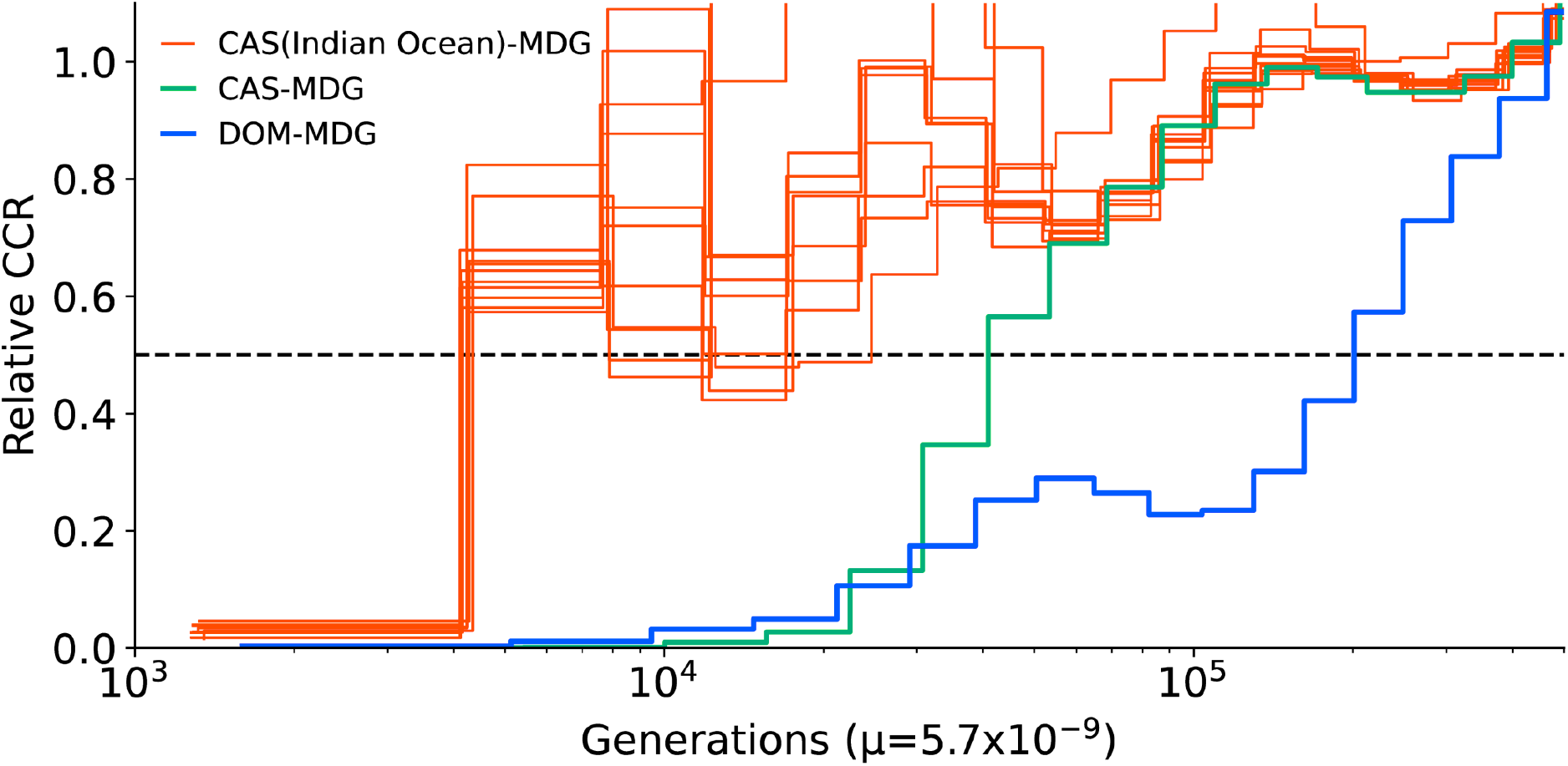
The figure shows the relative cross coalescent rate (rCCR) between the CAS(Indian Ocean)-MDG, CAS-MDG, and DOM-MDG inter-populations. The *x*-axis represents generations before the present scaled by a mutation rate of 0.57 × 10^−8^ per site per generation. The rCCR=0.5 is assumed heuristically as the time for the two populations to separate.

### The genomic pattern of CAS/DOM admixture in the Madagascar samples

In order to understand the genomic pattern of admixture in the Madagascar samples, we performed local ancestry estimation of the Madagascar samples, using the Indian CAS and German DOM as source populations (Supplementary Fig. S6). The method infers whether particular genomic regions of each Madagascar sample are derived from CAS or DOM. We confirmed that the genomes of Madagascar samples were mosaics of CAS and DOM (approximately 61% CAS ancestry), which is consistent with the results of PCA/Admixture/*f*_4_-statistics/MCMC in this study. Some of the DOM fragments were very long, up to about 40 Mbp. Such long DOM fragments are thought to be the result of relatively recent DOM hybridization events. When we performed the same local ancestry estimation for the Indonesian samples, the Bali sample showed a barcode-like pattern and did not have long DOM fragments like the Madagascar samples. As for the Java sample, there were some long DOM fragments, but they were not as long as the tracts in the Madagascar samples (Supplementary Fig. S7). These results indicate that the Madagascar samples experienced recent admixture with DOM.

## Discussion

The house mouse is an animal that has now colonized a wide range of areas including remote islands and archipelagos. This creature has been in a commensal with humans for at least 10,000 years (Cucchi *et al*., 2005), spread to Europe thousands of years ago (Jones *et al*., 2011), and is thought to have been introduced to remote islands, including Madagascar, by human economic activities over thousands or hundreds of years. Other than Madagascar, New Zealand (Searle *et al*., 2009a), Madeira (Gündüz *et al*., 2001), and the British Isles (Searle *et al*., 2009b) are known for their studies of the introduction of house mice to the island. In this study, we will provide insight into what subspecies of mice inhabiting the remote island of Madagascar belongs to and what genetic diversity they possess. However, we should make a caveat that this study was conducted using a limited number of samples captured in a small region, and a large-scale sampling across the island would benefit to draw a clearer picture in the future.

### Genetic background of the Madagascar wild house mouse samples

Previous studies have shown that the wild *M. musculus* exhibit a very large effective population size and nucleotide diversity compared to humans. The Madagascar house mouse samples analyzed in this study also showed a higher nucleotide diversity of 0.31% compared with humans (0.08%–0.12%) (Arbiza *et al*., 2014; Prado-Martinez *et al*., 2013; Perry *et al*., 2012); however, this value is less than half of the nucleotide diversity in CAS (0.74%–0.79%), slightly higher than the nucleotide diversity in MUS (0.18%–0.25%), and within the range of the nucleotide diversity in DOM (0.07%–0.35%) (Harr *et al*., 2016; Geraldes *et al*., 2008; Phifer-Rixey *et al*., 2014). Given that the samples used in this study were captured in the inland of Madagascar, the relatively low nucleotide diversity within the Madagascar wild house mouse samples may be related to the fact that these Madagascar samples either experienced bottlenecks when they were first brought to the island or were brought from multiple origins but their genetic features were similar.

Based on the results of the PCA plot, ADMIXTURE, *f*_4_-statistics test, phylogenetic tree, and phylogenetic network, we conclude that the nuclear genomes of Madagascar samples largely consisted of a CAS genetic component. This result is consistent with a previous microsatellite study (Hardouin *et al*., 2015). However, PCA, ADMIXTURE, *f*_4_-statistic test, and MSMC analysis demonstrated that our Madagascar samples have experienced admixture events with DOM. In addition, the local ancestry analysis showed long tracts with DOM genetic components in some samples, suggesting that an admixture is an ongoing event. The recent quantitative trait loci study of genes associated with hybrid fitness of *M. musculus* identified multiple regions causing hybrid male sterility in CAS and DOM hybrids (White *et al*., 2012). According to the study, a fairly high percentage of male individuals in the hybrid second generation (*F*_2_), exhibited phenotypes associated with infertility, indicating that CAS and DOM hybrids have difficulty in constructing long-lasting hybrids. Nevertheless, the Madagascar samples exhibit the DOM-like CAS genetic feature, indicating that inter-subspecific hybridization would be rare but possible in Madagascar.

The mitochondrial genome analysis indicates that the mitochondrial lineages of the Madagascar samples consist of a monophyletic group with low nucleotide diversity and that these lineages have a recent and unique origin with the tMRCA of approximately 3,000 years ago. MSMC analysis using autosomal haplotypes of Madagascar showed that our Madagascar samples experienced a bottleneck event about 1000–3000 years ago. This result is consistent with the mitochondrial analysis and suggests that *M. musculus* may have been brought to the island of Madagascar during this period. These patterns support the previous mitochondrial DNA study by Duplantier et al. (Duplantier *et al*., 2002; Sakuma *et al*., 2016). However, the study by Duplantier et al. (2002), which analyzed 539 nucleotide sites in the D-Loop of the mitochondrial genome, differs from our results, where the MUS clade is located at the basal of all *M. musculus*. We created a phylogenetic tree with approximately 16,000 bp and showed the mitochondrial genealogy with higher resolution than previous analyses, however, it should be noted that the bootstrap value supporting the basal position of Madagascar mitochondrial lineage was 0.64, which is not sufficiently high to conclude. As a note, the bootstrap values were equally low for both the previous partial mitochondrial sequence results and the current complete mitochondrial sequence results.

Collectively, our study showed highly complex genetic features of Madagascar samples. The nuclear genomes are mostly derived from CAS, but a significant amount of admixture with DOM was detected as well. In contrast, the mitochondrial lineage of Madagascar samples has highly diverged from both CAS and DOM lineage. These findings indicate that the genetic background of Madagascar house mice is highly unique among the other worldwide populations.

### Divergence of Madagascar samples from other CAS samples

We examined which CAS samples are the most closely related to the Madagascar samples. Although the neighbor-joining tree and neighbor-net network showed that the Madagascar samples were clustered with Indian and Nepalese samples, *f*_3_ statistics and MCMC analysis showed that the CAS samples in a wide range of the coastal and island regions of the Indian Ocean are genetically close to the Madagascar samples than the inland samples. Considering the fact that the inclusion of highly admixed samples to the tree and network constructions resulted in a distorted pattern of subspecies divergence, we concluded that the results of *f*_*3*_ statistics and MCMC would be more reliable. In particular, the model of MSMC assumes the coalescent time of genomic fragments between chromosomes differ across the genome and the estimated divergence time would be more robust to gene flow and admixture events. The relatively recent divergence time (∼4000 years ago) between Madagascar and the Indian Ocean coastal populations suggests the migration of the Madagascar population was related to human activity in the Neolithic period.

Notably, there was a small but clear increase of rCCR between the DOM and Madagascar populations around 60,000–100,000 years ago (Fig. 7). This pattern would be due to an admixture event between the ancestral population of German DOM and Madagascar. However, we should note that the peak timing (60,000–100,000 years ago) does not necessarily represents the actual time of the admixture event. Because we used German samples as representatives of DOM, if unsampled or extinct DOM population is an actual source population of admixture, the peak timing would rather represent the divergence time between the ancestral populations of German DOM and the unsampled/extinct DOM. Unfortunately, our dataset did not cover a large number of DOM samples, and we were not able to examine the timing of historical admixture events if existed. Caution should be exercised with these PSMC and MSMC results, especially for PSMC population size change peaks, as the apparent effective population size changes with migration due to population structure effects, and this may not reflect the true population history (Mazet et al., 2016).

### Possible history of the Madagascar wild house mouse samples

The ancestors of present Madagascar people are thought to have dual origins from the Indonesian island of Borneo in Southeast Asia and the eastern coast of Africa, and it is thought that wild house mouse was introduced to Madagascar in a commensal manner with the migration of these human ancestors who arrived on the Madagascar island.

We first consider when the ancestors of the Madagascar house mouse were introduced to the island. Although the molecular evolution rate of mitochondria, the mutation rate in nuclear genomes, and the generation time of mice are debatable issues and estimated time would be highly vulnerable with these assumptions, our mitochondrial genealogy and MSMC analysis consistently showed that they experienced a strong population bottleneck around 1000–3000 years ago. Although the estimated time range is wide, it is equivalent to the commonly accepted timing of migration when Austronesian-speaking people from the Indonesian islands arrived in Madagascar (BCE 300–CE 500) (Dewar and Wright, 1993; Burney *et al*., 2004), as well as the estimated timing using human genetic data (Hurles *et al*., 2005). Furthermore, cross-population MSMC analyses showed that the divergence time between the Madagascar samples and the coastal and island regions of South and Southeast Asia was approximately 4000 years ago. Since commensal animals, such as house mice, would not suddenly expand their population size without a proper type of human activity, such as agriculture, we suggest that the Madagascar house mice migrated to the island simultaneously or soon after the first Austronesian-speaking farmers arrived on the island. As for methods of divergence time estimation, mitochondrial molecular clocks have been well studied, but in this study, we estimated divergence time from various perspectives using not only mitochondria but also nuclear genomes (see Methods for more detail). It is important to note that these estimates are strongly dependent on the mutation rates and generation times used, which can double or halve the results.

The question of where the ancestors of Madagascar samples were migrated from is difficult to answer. Our *f*_3_ and MSMC studies showed that the Madagascar samples have a genetic affinity to many samples from coastal and island regions of the Indian Ocean and the origin was hard to be specified. In addition, our samples did not cover Yemen, where previous mitochondrial studies suggest the origin, and Borneo Island, where Austronesian-speaking people migrated from. The homeland of original Madagascar samples remains an open question.

Whether the introduction of house mice to Madagascar occurred multiple times or not is also an interesting question. Previous mitochondrial analysis on rats collected from multiple sites in Madagascar suggested that there was considerable genetic diversity among sites even within Madagascar, supporting that they originated from multiple places (Tollenaere *et al*., 2010). In this study, we showed that the Madagascar house mice have both CAS and DOM genetic features. The signature of introgression from DOM is absent in the Southeast Asian samples except for the Indonesian samples. The pattern of very recent introgression in Madagascar was qualitatively different from those in Indonesian samples, suggesting that the introgression from DOM, at least for the most recent one, has occurred in Madagascar, supporting multiple origins. A clearer view of the house mouse may emerge if samples could be collected from multiple sites throughout Madagascar.

## Conclusion

This study determined the whole-genome sequences of five Madagascar wild house mice and analyzed their genetic backgrounds. Until today, mice from Madagascar have been treated as DOM or *M. m. gentilulus*, but genomic evidence indicates that their genetic feature is mostly derived from CAS with a non-negligible amount of introgression from DOM. In addition, mitochondrial and nuclear genome evidence indicates that they experienced a bottleneck approximately 1000–3000 years ago, coinciding with the arrival of Austronesian humans to Madagascar. In the future, mouse samples from the Middle East and Borneo, as well as more samples across Madagascar, would clarify the genetic backgrounds of the Madagascar mouse population. Overall, this study revealed unique and highly complex genetic features of Madagascar house mice and provides insight into the rodents that have successfully migrated to Madagascar, which has only been partially understood.

## Data Archiving

The short-read whole-genome sequencing data generated in this study have been submitted to the DDBJ BioProject database (https://www.ddbj.nig.ac.jp/bioproject/) under the accession number PRJDB11969. The complete mitochondrial genome sequences are submitted to the DDBJ database under the accession number LC644158–LC644162.

## Acknowledgements

This work was partly supported by the MEXT KAKENHI (grant 18H05511 to N. O. and grant 18H05508 to H. S.). We would like to express my gratitude to Dr. Chihiro Tanaka, who belongs to the Animal Care and Exhibition Section, Yagiyama Zoological Park, Sendai, Japan, for her support of the sample collection project. We would also like to express my deepest gratitude to Drs. Kimiyuki Tsuchiya and Hajanirina Ramino for sample collection in Madagascar.

