## Supplementary Fig. for "Whole-genome sequencing analysis of wild-caught house mice *Mus musculus* from Madagascar"

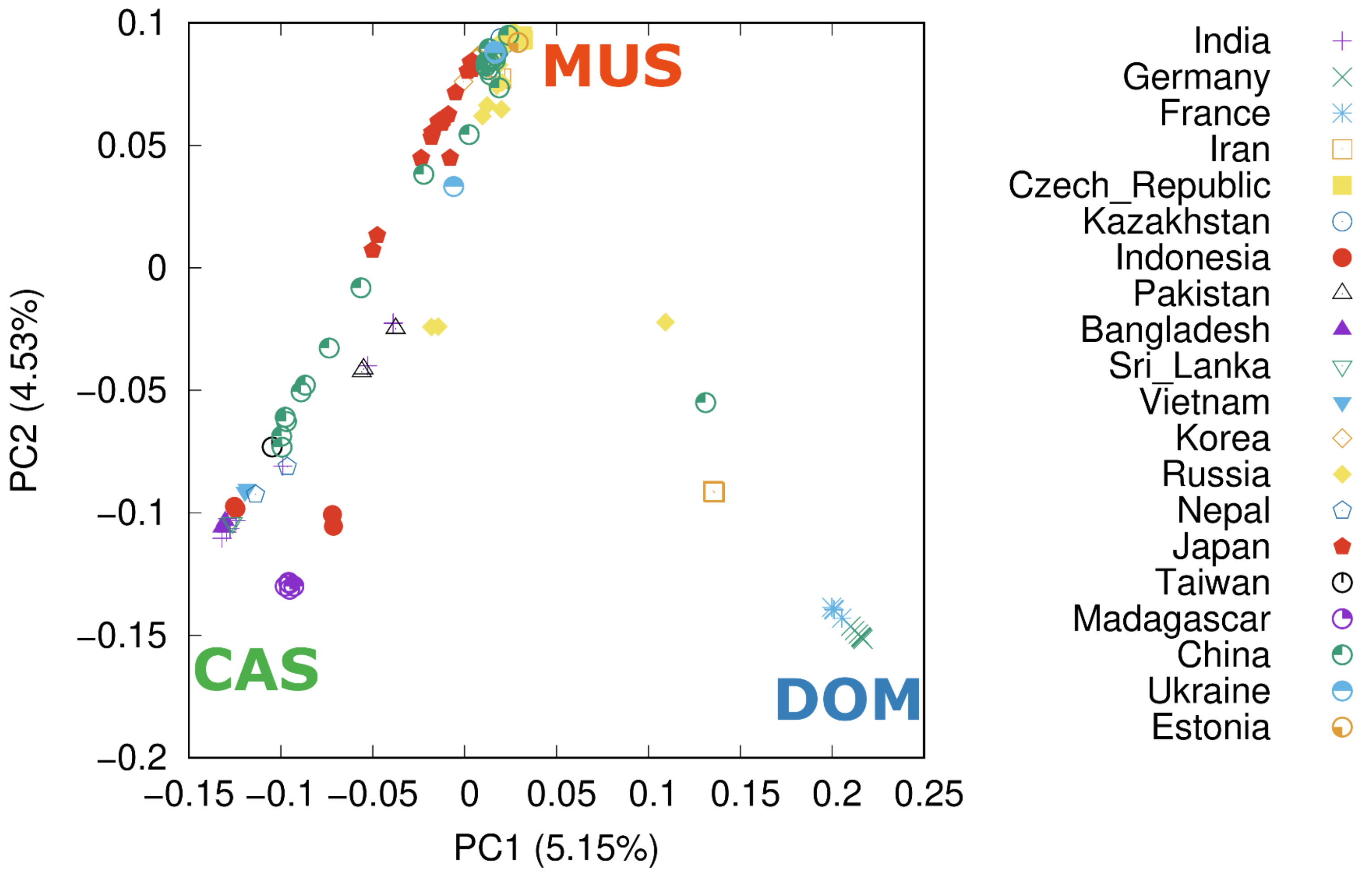


**Supplementary Fig. S1**

Principal Component Analysis plot of *M. musculus* using LD pruned autosomal single nucleotide variants (SNVs). The points in the figure represent the eigenvalues of individual samples collected from the countries shown on the right. The upper left, lower left, and right vertices of the triangle represent *Mus musculus domesticus* (DOM), *M. m. castaneus* (CAS), and *M. m. musculus* (MUS) genetic components, respectively. The proportion of variance for each eigenvalue is shown in parentheses on the labels of the *x*-axis and *y*-axis.

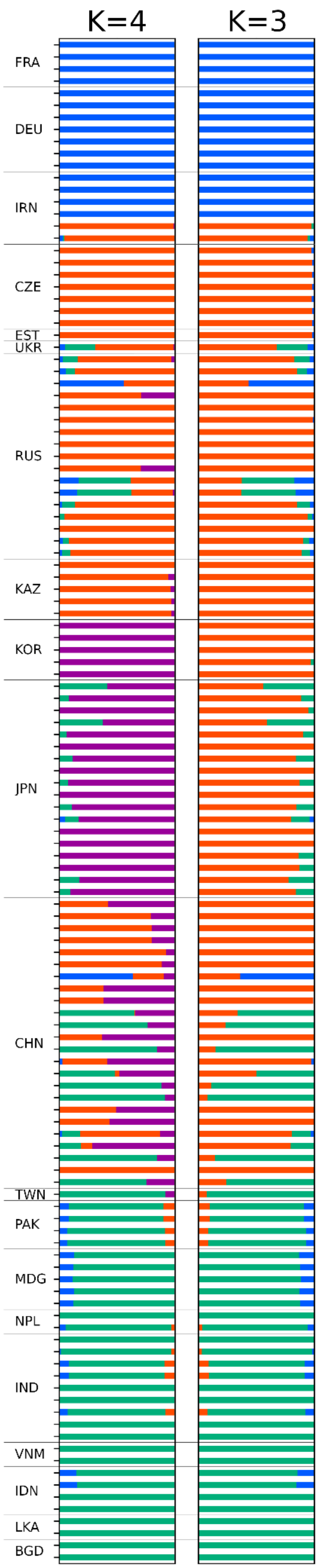


**Supplementary Fig. S2**

ADMIXTURE plot of autosomes. The plot shows the proportion of estimated subspecies' genetic components. Results for cluster $K$ = 3 and $K$ = 4 are presented. Samples with the same country code are ordered according to sampling site from west to east and north to south.

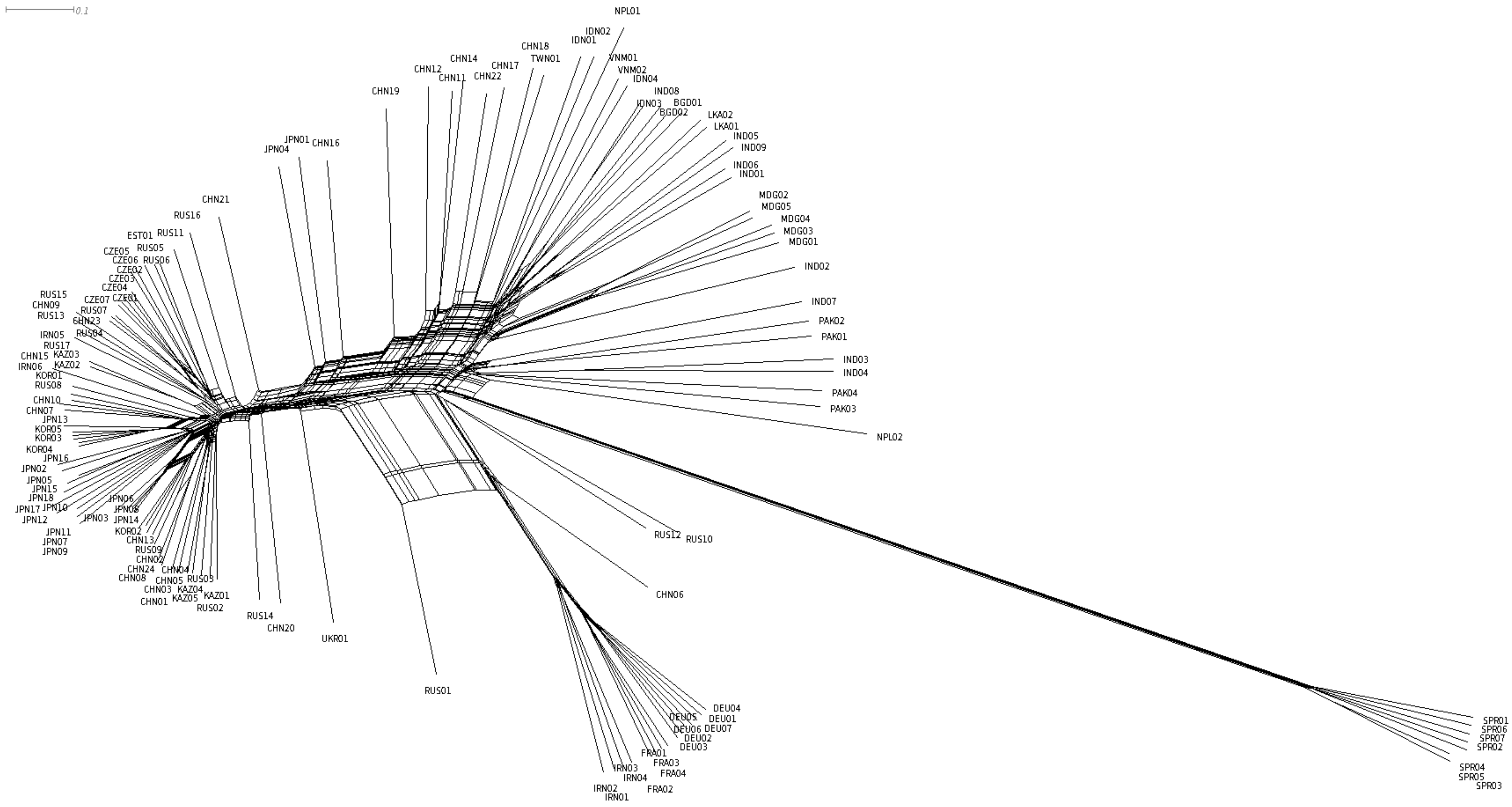


**Supplementary Fig. S3**

The Neighbor-Net network was constructed with the whole genome genetic distance using all samples in this study. The labels are based on the sample code described in Supplementary Table 1.

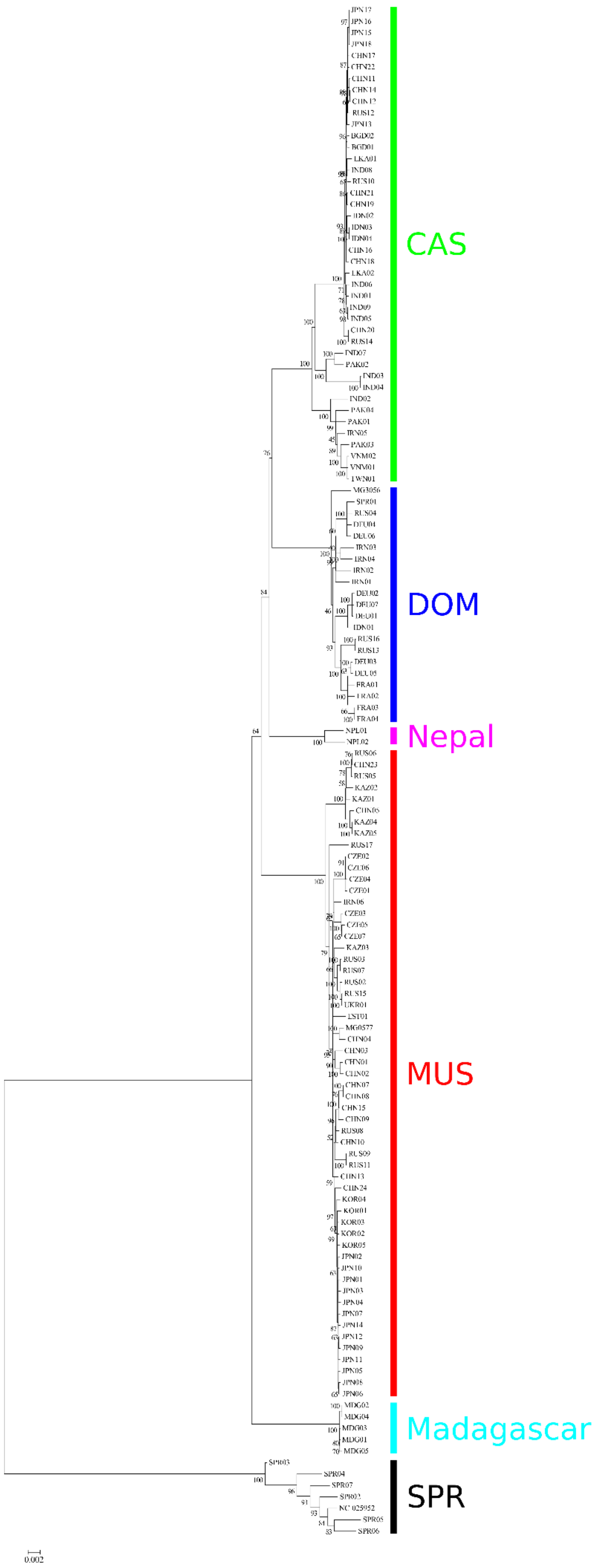


**Supplementary Fig. S4**

The maximum likelihood tree using mitochondrial genomes. The substitution model (TIM2+F+R3) was determined by the best fit model of ModelFinder implemented in IQ-TREE. The colored line and captions correspond to the mitochondrial genome clades of *Mus musculus*. The numbers exhibit the results of the bootstrap resampling with 1000 replications.

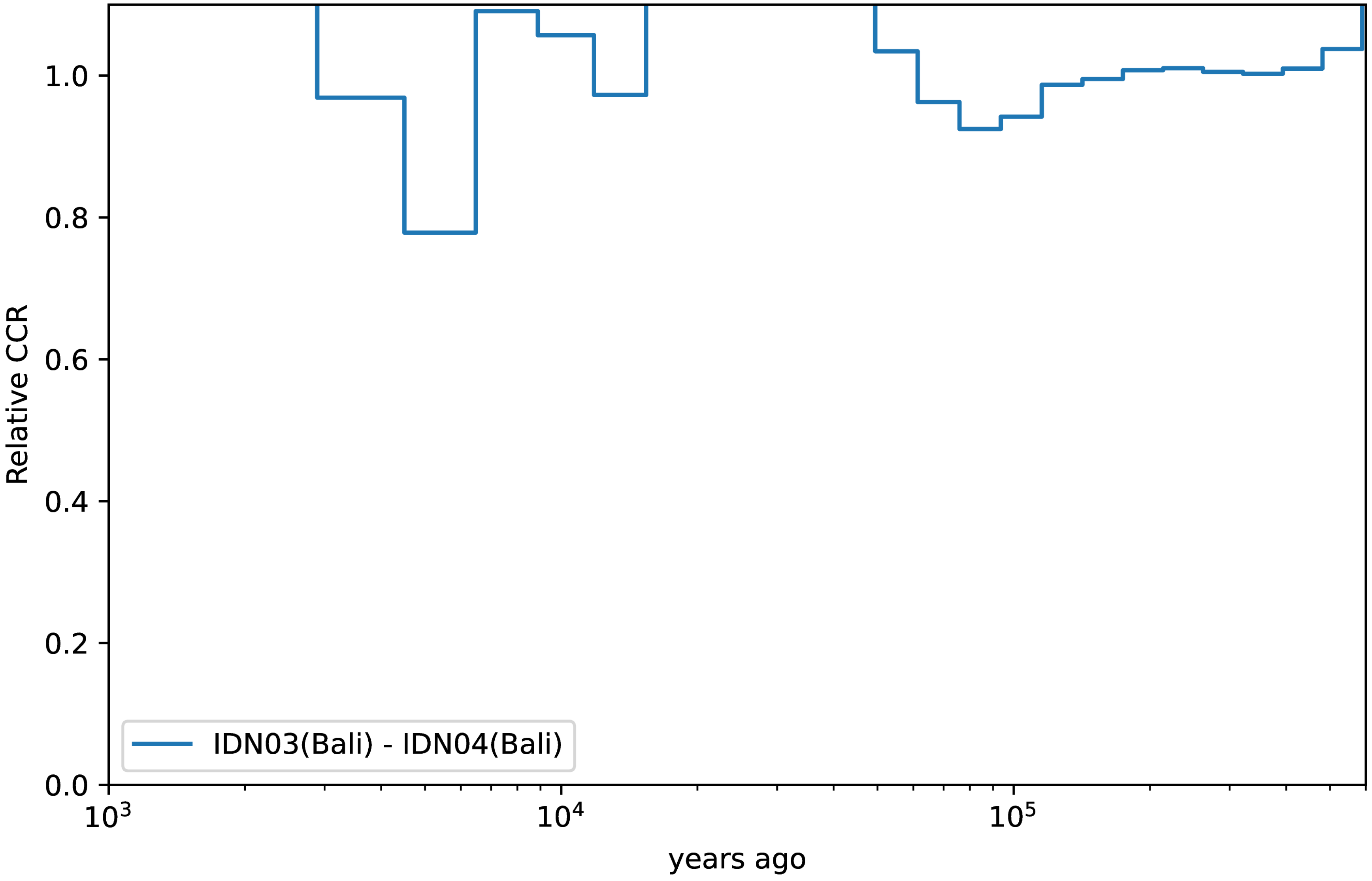


**Supplementary Fig. S5**
The rCCR plot of two Bali, Indonesia samples as control. If the two samples were divergent, the rCCR should show less than 0.5, but since these two samples are from the same population, they do not show such a trend.

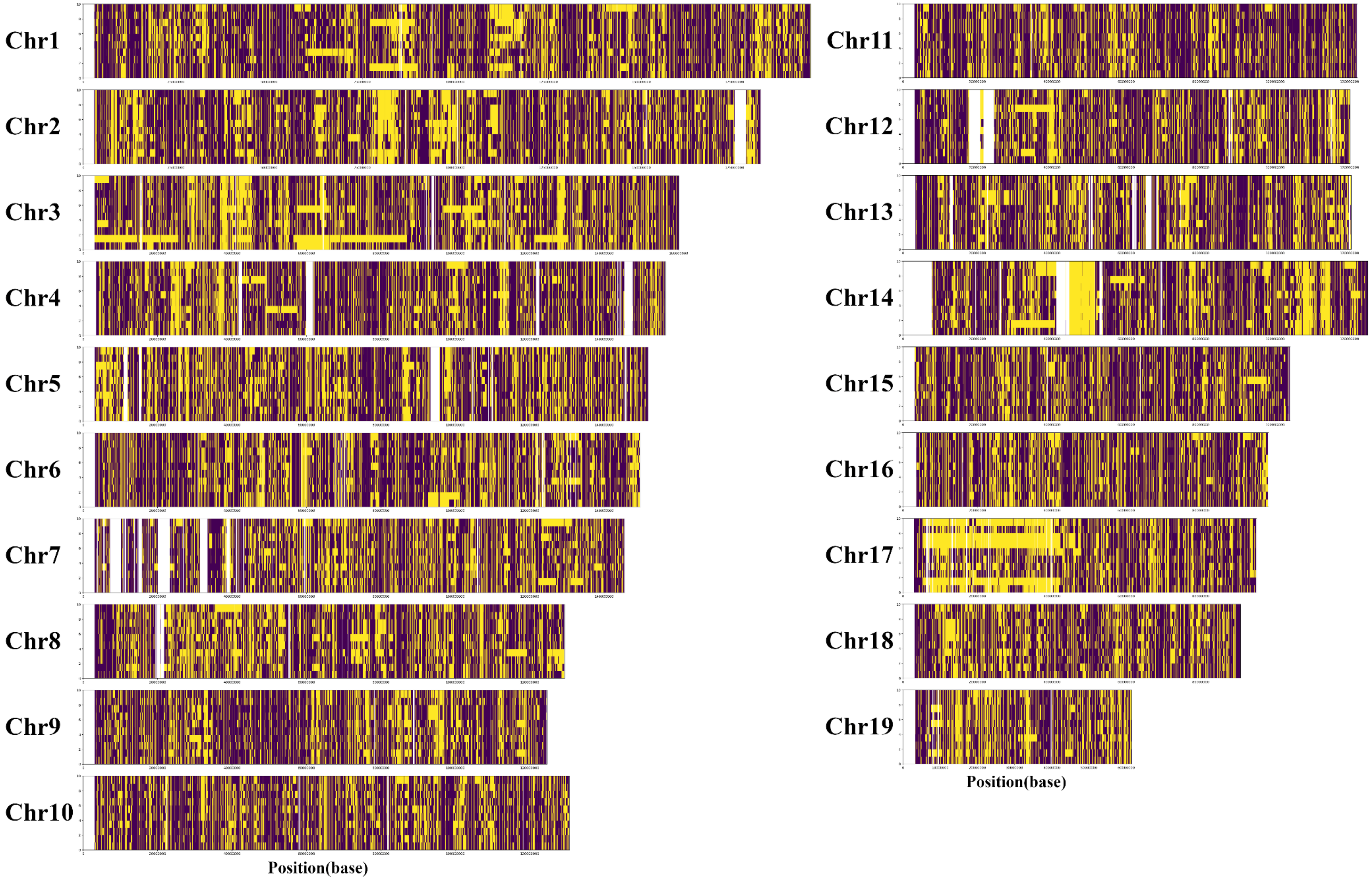


**Supplementary Fig. S6**

The local ancestry inference plot of Madagascar samples. The DOM samples from Germany and CAS samples from India were used as reference donor populations. The *x*-axis of each chromosome shows the haplotypes (10 haplotypes from five samples). The *y*-axis of each chromosome shows the position. The yellow fragments show the DOM-derived SNPs and the purple fragments show the CAS-derived SNPs. Blank areas represent positions where SNPs were not detected (or not mapped).

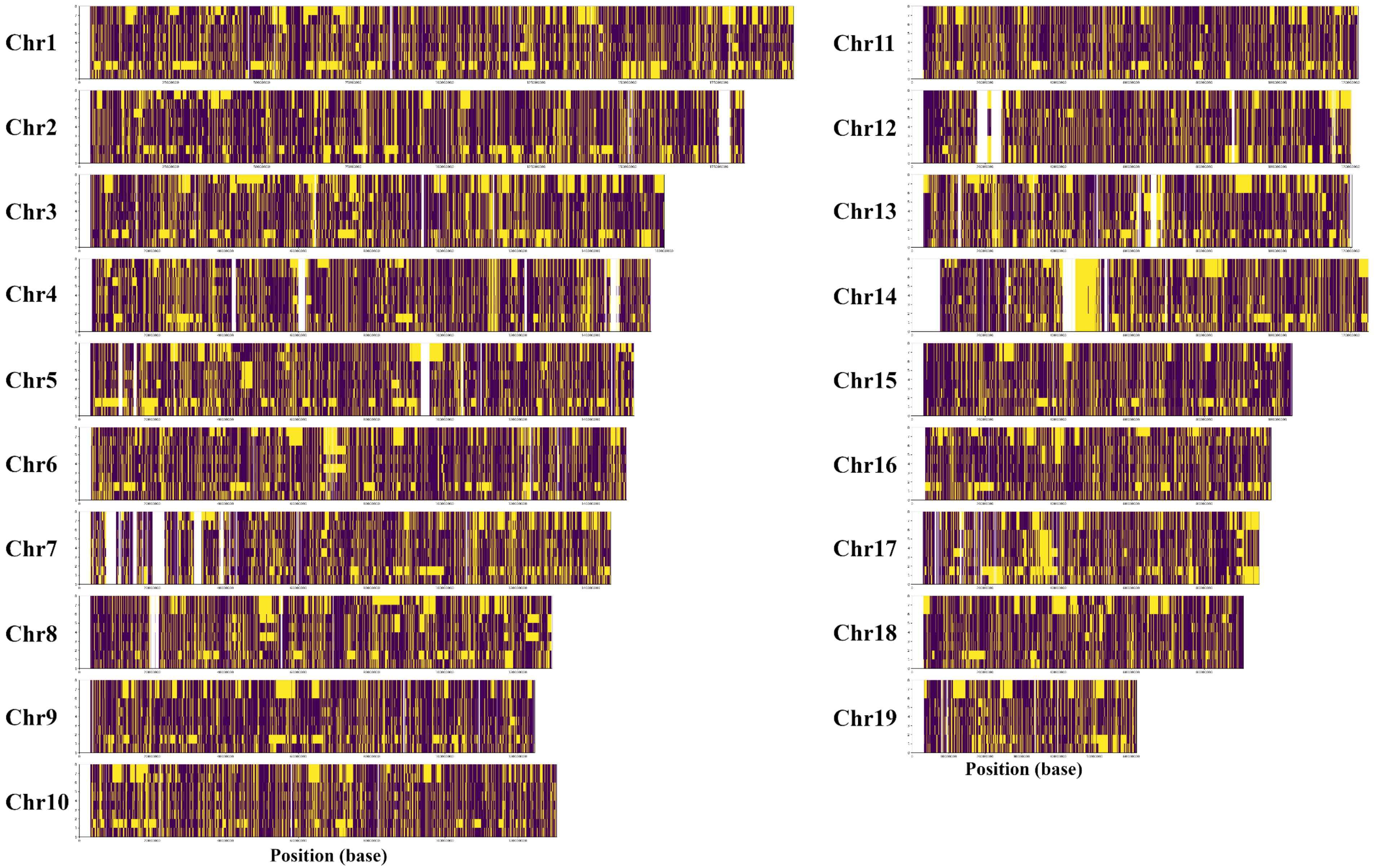


**Supplementary Fig. S7**

The local ancestry inference plot of Indonesian samples. The DOM samples from Germany and CAS samples from India were used as reference donor populations. The *x*-axis of each chromosome shows the haplotypes (8 haplotypes from four samples). The *y*-axis of each chromosome shows the position. The yellow fragments show the DOM-derived SNPs and the purple fragments show the CAS-derived SNPs. Blank areas represent positions where SNPs were not detected (or not mapped).
