## Supplementary Table 1 for "Whole-genome sequencing analysis of wild-caught house mice *Mus musculus* from Madagascar"

**Supplementary Table 1. Sample list of whole genome sequencing data of wild house mouse used in this study.**

| Sample Code | Sample ID | mtDNA Haplogroup | Location | **Latitude | **Longitude | Reference |
|---|---|---|---|---|---|---|
| BGD01 | HI0373 | CAS | Bangladesh: Dhaka | 23.702000 | 90.366944 | Fujiwara et al. 2021, Li et al. 2021 |
| BGD02 | HI0347 | CAS | Bangladesh: Mymensingh | 24.753889 | 90.403056 | Fujiwara et al. 2021, Li et al. 2021 |
| CHN01 | MG0686 | MUS | China: Kashgar | 39.468100 | 75.993800 | Fujiwara et al. 2021, Li et al. 2021 |
| CHN02 | MG0747 | MUS | China: Hotan | 37.100000 | 80.016667 | Fujiwara et al. 2021, Li et al. 2021 |
| CHN03 | MG0608 | MUS | China: Aksu | 41.185000 | 80.290400 | Fujiwara et al. 2021, Li et al. 2021 |
| CHN04 | MG0611 | MUS | China: Tacheng | 46.751700 | 82.986900 | Fujiwara et al. 2021, Li et al. 2021 |
| CHN05 | MG0597 | MUS | China: Manas | 44.300000 | 86.216667 | Fujiwara et al. 2021, Li et al. 2021 |
| CHN06 | MG0577 | MUS | China: Urumqi | 43.800000 | 87.583333 | Fujiwara et al. 2021, Li et al. 2021 |
| CHN07 | MG0716 | MUS | China: Lhasa | 29.653400 | 91.171900 | Fujiwara et al. 2021, Li et al. 2021 |
| CHN08 | MG0721 | MUS | China: Lhasa | 29.653400 | 91.171900 | Fujiwara et al. 2021, Li et al. 2021 |
| CHN09 | MG0871 | MUS | China: Dunhuang | 40.142100 | 94.661900 | Fujiwara et al. 2021, Li et al. 2021 |
| CHN10 | MG5086 | MUS | China: Jiayuguan | 39.773200 | 98.288200 | Fujiwara et al. 2021, Li et al. 2021 |
| CHN11 | MG0917 | CAS | China: Lijiang | 26.855200 | 100.225900 | Fujiwara et al. 2021, Li et al. 2021 |
| CHN12 | MG0797 | CAS | China: Dali | 25.606500 | 100.267600 | Fujiwara et al. 2021, Li et al. 2021 |
| CHN13 | MG0565 | MUS | China: Xining | 36.616667 | 101.766667 | Fujiwara et al. 2021, Li et al. 2021 |
| CHN14 | MG0529 | CAS | China: Kunming | 25.046400 | 102.709400 | Fujiwara et al. 2021, Li et al. 2021 |
| CHN15 | MG0507 | MUS | China: Lanzhou | 36.060600 | 103.826800 | Fujiwara et al. 2021, Li et al. 2021 |
| CHN16 | MG0709 | CAS | China: Chongqing | 29.563700 | 106.550400 | Fujiwara et al. 2021, Li et al. 2021 |
| CHN17 | MG0501 | CAS | China: Guilin | 25.275000 | 110.296000 | Fujiwara et al. 2021, Li et al. 2021 |
| CHN18 | MG0504 | CAS | China: Guangzhou | 23.132000 | 113.266000 | Fujiwara et al. 2021, Li et al. 2021 |
| CHN19 | MG0908 | CAS | China: Wuhan | 30.593400 | 114.304600 | Fujiwara et al. 2021, Li et al. 2021 |
| CHN20 | MG0863 | CAS | China: Manzhouli | 49.598000 | 117.379000 | Fujiwara et al. 2021, Li et al. 2021 |
| CHN21 | MG0713 | CAS | China: Zhenjiang | 32.188000 | 119.424000 | Fujiwara et al. 2021, Li et al. 2021 |
| CHN22 | MG0795 | CAS | China: Ningbo | 29.860300 | 121.624500 | Fujiwara et al. 2021, Li et al. 2021 |
| CHN23 | MG0631 | MUS | China: Mohe | 52.972000 | 122.539000 | Fujiwara et al. 2021, Li et al. 2021 |
| CHN24 | MG0992 | MUS | China: Qiqihar | 47.354900 | 123.918200 | Fujiwara et al. 2021, Li et al. 2021 |
| CZE01 | ERS957497 | MUS | Czech Republic: Studenec | see Harr et al. 2016 | | Harr et al. 2016 |
| CZE02 | ERS957498 | MUS | Czech Republic: Studenec | see Harr et al. 2016 | | Harr et al. 2016 |
| CZE03 | ERS957499 | MUS | Czech Republic: Studenec | see Harr et al. 2016 | | Harr et al. 2016 |
| CZE04 | ERS957500 | MUS | Czech Republic: Studenec | see Harr et al. 2016 | | Harr et al. 2016 |
| CZE05 | ERS957501 | MUS | Czech Republic: Studenec | see Harr et al. 2016 | | Harr et al. 2016 |
| CZE06 | ERS957502 | MUS | Czech Republic: Studenec | see Harr et al. 2016 | | Harr et al. 2016 |
| CZE07 | ERS957503 | MUS | Czech Republic: Studenec | see Harr et al. 2016 | | Harr et al. 2016 |
| DEU01 | ERS739373 | DOM | Germany: Cologne-Bonn | see Harr et al. 2016 | | Harr et al. 2016 |
| DEU02 | ERS739374 | DOM | Germany: Cologne-Bonn | see Harr et al. 2016 | | Harr et al. 2016 |
| DEU03 | ERS739375 | DOM | Germany: Cologne-Bonn | see Harr et al. 2016 | | Harr et al. 2016 |
| DEU04 | ERS739376 | DOM | Germany: Cologne-Bonn | see Harr et al. 2016 | | Harr et al. 2016 |
| DEU05 | ERS739377 | DOM | Germany: Cologne-Bonn | see Harr et al. 2016 | | Harr et al. 2016 |
| DEU06 | ERS739378 | DOM | Germany: Cologne-Bonn | see Harr et al. 2016 | | Harr et al. 2016 |
| DEU07 | ERS739379 | DOM | Germany: Cologne-Bonn | see Harr et al. 2016 | | Harr et al. 2016 |
| EST01 | MG3044 | MUS | Estonia: Tallinn | 59.437222 | 24.745278 | Fujiwara et al. 2021, Li et al. 2021 |
| FRA01 | ERS739382 | DOM | France: Massif Central | see Harr et al. 2016 | | Harr et al. 2016 |
| FRA02 | ERS739385 | DOM | France: Massif Central | see Harr et al. 2016 | | Harr et al. 2016 |
| FRA03 | ERS739386 | DOM | France: Massif Central | see Harr et al. 2016 | | Harr et al. 2016 |
| FRA04 | ERS739388 | DOM | France: Massif Central | see Harr et al. 2016 | | Harr et al. 2016 |
| IDN01 | HI0112 | DOM | Indonesia: Bogor | -6.600000 | 106.800000 | Fujiwara et al. 2021, Li et al. 2021 |
| IDN02 | HI0134 | CAS | Indonesia: Lembang | -6.817000 | 107.617000 | Fujiwara et al. 2021, Li et al. 2021 |
| IDN03 | HI0115 | CAS | Indonesia: Bali | -8.369167 | 115.138333 | Fujiwara et al. 2021, Li et al. 2021 |
| IDN04 | HI0116 | CAS | Indonesia: Bali | -8.369167 | 115.138333 | Fujiwara et al. 2021, Li et al. 2021 |
| IND01 | HI0274 | CAS | India: Mysore | 12.308611 | 76.653056 | Fujiwara et al. 2021, Li et al. 2021 |
| IND02 | HI0185 | CAS | India: Delhi | 28.610000 | 77.230000 | Fujiwara et al. 2021, Li et al. 2021 |
| IND03 | HI0159 | CAS | India: Leh | 34.160000 | 77.580000 | Fujiwara et al. 2021, Li et al. 2021 |
| IND04 | HI0161 | CAS | India: Leh | 34.160000 | 77.580000 | Fujiwara et al. 2021, Li et al. 2021 |
| IND05 | HI0328 | CAS | India: Hyderabad | 17.370000 | 78.480000 | Fujiwara et al. 2021, Li et al. 2021 |
| IND06 | HI0329 | CAS | India: Hyderabad | 17.370000 | 78.480000 | Fujiwara et al. 2021, Li et al. 2021 |
| IND07 | ERS003044 | CAS | India: Himalaya | see Harr et al. 2016 | | Harr et al. 2016 |
| IND08 | HI0313 | CAS | India: Bhubaneswar | 20.270000 | 85.840000 | Fujiwara et al. 2021, Li et al. 2021 |
| IND09 | HI0321 | CAS | India: Bhubaneswar | 20.270000 | 85.840000 | Fujiwara et al. 2021, Li et al. 2021 |
| IRN01 | ERS739389 | DOM | Iran: Ahvaz | see Harr et al. 2016 | | Harr et al. 2016 |
| IRN02 | ERS739393 | DOM | Iran: Ahvaz | see Harr et al. 2016 | | Harr et al. 2016 |
| IRN03 | ERS739395 | DOM | Iran: Ahvaz | see Harr et al. 2016 | | Harr et al. 2016 |
| IRN04 | ERS739396 | DOM | Iran: Ahvaz | see Harr et al. 2016 | | Harr et al. 2016 |
| IRN05 | MG5135 | CAS | Iran: Now Shahr | 36.648889 | 51.496111 | Fujiwara et al. 2021, Li et al. 2021 |
| IRN06 | MG0417 | MUS | Iran: Mashhad | 36.300000 | 59.600000 | Fujiwara et al. 2021, Li et al. 2021 |
| JPN01 | HS2788 | MUS | Japan: Okinawa Island | 26.212444 | 127.680917 | Fujiwara et al. 2021, Li et al. 2021 |
| JPN02 | HS4272 | MUS | Japan: Fukuoka | 33.590139 | 130.401722 | Fujiwara et al. 2021, Li et al. 2021 |
| JPN03 | HS4273 | MUS | Japan: Yamaguchi | 34.178028 | 131.473778 | Fujiwara et al. 2021, Li et al. 2021 |
| JPN04 | HS4411 | MUS | Japan: Oda | 35.192083 | 132.499361 | Fujiwara et al. 2021, Li et al. 2021 |
| JPN05 | HS4097 | MUS | Japan: Misasa | 35.408444 | 133.861778 | Fujiwara et al. 2021, Li et al. 2021 |

| | | | | | | |
|---|---|---|---|---|---|---|
| JPN06 | HS4120 | MUS | Japan: Ishii | 34.074167 | 134.440556 | Fujiwara et al. 2021, Li et al. 2021 |
| JPN07 | HS2815 | MUS | Japan: Nonoichi | 36.519417 | 136.609778 | Fujiwara et al. 2021, Li et al. 2021 |
| JPN08 | HS4056 | MUS | Japan: Fujisawa | 35.338833 | 139.491111 | Fujiwara et al. 2021, Li et al. 2021 |
| JPN09 | HS4271 | MUS | Japan: Setana | 42.417111 | 139.883028 | Fujiwara et al. 2021, Li et al. 2021 |
| JPN10 | HS4169 | MUS | Japan: Tsukuba | 36.083472 | 140.076444 | Fujiwara et al. 2021, Li et al. 2021 |
| JPN11 | HS3534 | MUS | Japan: Yamagata | 38.255417 | 140.339611 | Fujiwara et al. 2021, Li et al. 2021 |
| JPN12 | HS2323 | MUS | Japan: Hakodate | 41.768667 | 140.728917 | Fujiwara et al. 2021, Li et al. 2021 |
| JPN13 | MG0230 | CAS | Japan: Ashiro | 40.101389 | 141.052028 | Fujiwara et al. 2021, Li et al. 2021 |
| JPN14 | HS3538 | MUS | Japan: Tono | 39.702083 | 141.154500 | Fujiwara et al. 2021, Li et al. 2021 |
| JPN15 | MG0296 | CAS | Japan: Sapporo | 43.121917 | 141.245833 | Fujiwara et al. 2021, Li et al. 2021 |
| JPN16 | MG0297 | CAS | Japan: Sapporo | 43.121917 | 141.245833 | Fujiwara et al. 2021, Li et al. 2021 |
| JPN17 | HS2445 | CAS | Japan: Takikawa | 43.557833 | 141.910417 | Fujiwara et al. 2021, Li et al. 2021 |
| JPN18 | HS2326 | CAS | Japan: Nayoro | 44.355833 | 142.463333 | Fujiwara et al. 2021, Li et al. 2021 |
| KAZ01 | HS1464 | MUS | Kazakhstan: Aktobe | 50.283333 | 57.166667 | Fujiwara et al. 2021, Li et al. 2021 |
| KAZ02 | ERS957504 | MUS | Kazakhstan: Almaty | see Harr et al. 2016 | | Harr et al. 2016 |
| KAZ03 | ERS957506 | MUS | Kazakhstan: Almaty | see Harr et al. 2016 | | Harr et al. 2016 |
| KAZ04 | ERS957508 | MUS | Kazakhstan: Almaty | see Harr et al. 2016 | | Harr et al. 2016 |
| KAZ05 | ERS957510 | MUS | Kazakhstan: Almaty | see Harr et al. 2016 | | Harr et al. 2016 |
| KOR01 | HS0682 | MUS | South Korea: Baengnyeong Island | 37.966667 | 124.650000 | Fujiwara et al. 2021, Li et al. 2021 |
| KOR02 | HS4238 | MUS | South Korea: Ganghwa Island | 37.700000 | 126.433333 | Fujiwara et al. 2021, Li et al. 2021 |
| KOR03 | HS4233 | MUS | South Korea: Hwacheon-gun | 38.107500 | 127.714444 | Fujiwara et al. 2021, Li et al. 2021 |
| KOR04 | MG0444 | MUS | South Korea: Busan | 35.100000 | 129.033333 | Fujiwara et al. 2021, Li et al. 2021 |
| KOR05 | HS1368 | MUS | South Korea: Geoncheon | 35.837686 | 129.045187 | Fujiwara et al. 2021, Li et al. 2021 |
| LKA01 | HI0488 | CAS | Sri Lanka: Colombo | 6.934444 | 79.842778 | Fujiwara et al. 2021, Li et al. 2021 |
| LKA02 | HI0484 | CAS | Sri Lanka: Peradeniya | 7.266667 | 80.600000 | Fujiwara et al. 2021, Li et al. 2021 |
| MDG01 | HS5555 | GEN | Madagascar: Tsimbazaza | -18.93 | 47.526111 | Fujiwara et al. 2021, Li et al. 2021 |
| MDG02 | HS5558 | GEN | Madagascar: Tsimbazaza | -18.93 | 47.526111 | Fujiwara et al. 2021, Li et al. 2021 |
| MDG03 | HS5559 | GEN | Madagascar: Tsimbazaza | -18.93 | 47.526111 | Fujiwara et al. 2021, Li et al. 2021 |
| MDG04 | HS5560 | GEN | Madagascar: Tsimbazaza | -18.93 | 47.526111 | Fujiwara et al. 2021, Li et al. 2021 |
| MDG05 | HS5561 | GEN | Madagascar: Tsimbazaza | -18.93 | 47.526111 | Fujiwara et al. 2021, Li et al. 2021 |
| NPL01 | HS1467 | NEP | Nepal: Tukuche | 28.740000 | 83.620000 | Fujiwara et al. 2021, Li et al. 2021 |
| NPL02 | HS1523 | NEP | Nepal: Kathmandu | 27.717200 | 85.324000 | Fujiwara et al. 2021, Li et al. 2021 |
| PAK01 | HI0173 | CAS | Pakistan: Islamabad | 33.716667 | 73.066667 | Fujiwara et al. 2021, Li et al. 2021 |
| PAK02 | HI0175 | CAS | Pakistan: Islamabad | 33.716667 | 73.066667 | Fujiwara et al. 2021, Li et al. 2021 |
| PAK03 | HI0261 | CAS | Pakistan: Sahiwal | 30.661111 | 73.108333 | Fujiwara et al. 2021, Li et al. 2021 |
| PAK04 | HI0264 | CAS | Pakistan: Lahore | 31.549722 | 74.343611 | Fujiwara et al. 2021, Li et al. 2021 |
| RUS01 | MG3056 | DOM | Russia: Moscow | 55.755833 | 37.617222 | Fujiwara et al. 2021, Li et al. 2021 |
| RUS02 | MG3010 | MUS | Russia: North Caucasus | 43.711400 | 44.806100 | Fujiwara et al. 2021, Li et al. 2021 |
| RUS03 | HS3612 | MUS | Russia: Astrakhan | 46.350000 | 48.050000 | Fujiwara et al. 2021, Li et al. 2021 |
| RUS04 | HS3604 | DOM | Russia: Tomsk | 56.500000 | 84.966667 | Fujiwara et al. 2021, Li et al. 2021 |
| RUS05 | HS3605 | MUS | Russia: Gorno-Altaysk | 51.950000 | 85.966667 | Fujiwara et al. 2021, Li et al. 2021 |
| RUS06 | HS3608 | MUS | Russia: Irkutsk | 52.283333 | 104.283333 | Fujiwara et al. 2021, Li et al. 2021 |
| RUS07 | MG3012 | MUS | Russia: Chita | 52.050000 | 113.466667 | Fujiwara et al. 2021, Li et al. 2021 |
| RUS08 | MG3007 | MUS | Russia: Krementui | 50.155148 | 115.906841 | Fujiwara et al. 2021, Li et al. 2021 |
| RUS09 | MG3008 | MUS | Russia: Krementui | 50.155148 | 115.906841 | Fujiwara et al. 2021, Li et al. 2021 |
| RUS10 | HS1411 | CAS | Russia: Khasan | 42.428333 | 130.645556 | Fujiwara et al. 2021, Li et al. 2021 |
| RUS11 | MG3026 | MUS | Russia: Birakan | 49.000833 | 131.720000 | Fujiwara et al. 2021, Li et al. 2021 |
| RUS12 | MG3025 | CAS | Russia: Vladivostok | 43.133333 | 131.900000 | Fujiwara et al. 2021, Li et al. 2021 |
| RUS13 | MG3037 | DOM | Russia: Khabarovsk | 48.483333 | 135.083333 | Fujiwara et al. 2021, Li et al. 2021 |
| RUS14 | MG3018 | CAS | Russia: Rudnaya Pristan | 44.358611 | 135.810278 | Fujiwara et al. 2021, Li et al. 2021 |
| RUS15 | MG3048 | MUS | Russia: Yuzhno-Sakhalinsk | 46.966667 | 142.733333 | Fujiwara et al. 2021, Li et al. 2021 |
| RUS16 | HS3607 | DOM | Russia: Okha | 53.583333 | 142.933333 | Fujiwara et al. 2021, Li et al. 2021 |
| RUS17 | MG3045 | MUS | Russia: Poronaysk | 49.216667 | 143.116667 | Fujiwara et al. 2021, Li et al. 2021 |
| SPR01 | ERS957512 | *Mus spretus* | Spain: Madrid | see Harr et al. 2016 | | Harr et al. 2016 |
| SPR02 | ERS957513 | *Mus spretus* | Spain: Madrid | see Harr et al. 2016 | | Harr et al. 2016 |
| SPR03 | ERS957514 | *Mus spretus* | Spain: Madrid | see Harr et al. 2016 | | Harr et al. 2016 |
| SPR04 | ERS957515 | *Mus spretus* | Spain: Madrid | see Harr et al. 2016 | | Harr et al. 2016 |
| SPR05 | ERS957516 | *Mus spretus* | Spain: Madrid | see Harr et al. 2016 | | Harr et al. 2016 |
| SPR06 | ERS957517 | *Mus spretus* | Spain: Madrid | see Harr et al. 2016 | | Harr et al. 2016 |
| SPR07 | ERS957519 | *Mus spretus* | Spain: Madrid | see Harr et al. 2016 | | Harr et al. 2016 |
| TWN01 | HS2400 | CAS | Taiwan: Taitung | 22.758333 | 121.144444 | Fujiwara et al. 2021, Li et al. 2021 |
| UKR01 | MG3066 | MUS | Ukraine: Donetsk | 48.002778 | 37.805278 | Fujiwara et al. 2021, Li et al. 2021 |
| VNM01 | HI0505 | CAS | Viet Nam: Vinh Phuc | 21.300000 | 105.600000 | Fujiwara et al. 2021, Li et al. 2021 |
| VNM02 | HI0520 | CAS | Viet Nam: Hanoi | 21.028472 | 105.854167 | Fujiwara et al. 2021, Li et al. 2021 |

*Fujiwara K, Kawai Y, Moriwaki K, Takada T, Shiroishi T, Saitou N, Suzuki H, Osada N. 2021. Insights into Mus musculus subspecies population structure across Eurasia revealed by whole-genome sequence analysis. *Biorxiv* 2021.02.05.429881.

*Harr B, Karakoc E, Neme R, Teschke M, Pfeifle C, Pezer Ž, Babiker H, Linnenbrink M, Montero I, Scavetta R, et al. 2016. Genomic resources for wild populations of the house mouse, Mus musculus and its close relative Mus spretus. *Scientific Data* 3: 160075.

*Li Y, Fujiwara K, Osada N, Kawai Y, Takada T, Kryukov AP, Abe K, Yonekawa H, Shiroishi T, Moriwaki K, et al. 2020. House mouse Mus musculus dispersal in East Eurasia inferred from 98 newly determined complete mitochondrial genome sequences. *Heredity* 126: 132–147.

**Latitude and longitude are not absolute points of sample collection, but rather approximate locations.
