## Supplementary Table 2 for "Whole-genome sequencing analysis of wild-caught house mice *Mus musculus* from Madagascar"

**Supplementary Table 2. Sex of the collected sample by computing the depth of the sex chromosomes.**

| Sample Code | Sample ID | Chromosome X Total Depth | Chromosome Y Total Depth | Chromosome X (Normalized) | Chromosome Y (Normalized) | X/Y ratio | Sex |
|---|---|---|---|---|---|---|---|
| MDG01 | HS5555 | 1,659,509,030 | 15,751,830 | 18.324 | 17.654 | 1.038 | M |
| MDG02 | HS5558 | 3,180,178,155 | 116,836 | 35.114 | 0.131 | 268.160 | F |
| MDG03 | HS5559 | 1,704,627,115 | 15,290,145 | 18.822 | 17.136 | 1.098 | M |
| MDG04 | HS5560 | 1,686,612,760 | 15,456,122 | 18.623 | 17.323 | 1.075 | M |
| MDG05 | HS5561 | 3,089,337,937 | 205,312 | 34.111 | 0.230 | 148.242 | F |
