## Supplementary Table 3 for "Whole-genome sequencing analysis of wild-caught house mice *Mus musculus* from Madagascar"

**Supplementary Table 3. The $f_4$ statistics result of Madagascar wild house mouse population**

| Pop1(W) | Pop2(X) | Pop3(Y) | Pop4(Z) | F4-statistics | StdError | Z | BABA | ABBA | # of SNP |
|---|---|---|---|---|---|---|---|---|---|
| SPR | DOM | CAS | MDG | 0.006637 | 0.000253 | 26.25 | 2,030,967 | 1,348,330 | 102,858,288 |
| SPR | CAS | DOM | MDG | 0.005425 | 0.000103 | 52.426 | 1,906,338 | 1,348,330 | 102,858,288 |
| SPR | CAS | NEP | MDG | 0.005975 | 0.000213 | 28.091 | 2,021,931 | 1,407,379 | 102,858,288 |
| SPR | DOM | NEP | MDG | 0.011287 | 0.000251 | 44.953 | 2,310,098 | 1,149,146 | 102,858,288 |
